# Biological differences in promyelocytic leukemia (PML) proteins between PML-nuclear bodies (PML-NBs) and extranuclear PML bodies (EnPBs) in arsenite-exposed cells

**DOI:** 10.64898/2026.01.25.701640

**Authors:** Seishiro Hirano, Osamu Udagawa

**Affiliations:** Center for Health and Environmental Risk Research, National Institute for Environmental Studies, Japan

**Author notes:** Corresponding author: Seishiro Hirano, Ph.D., Center for Health and Environmental Risk Research, National Institute for Environmental Studies, 16-2 Onogawa, Tsukuba, Ibaraki 305-8506, Japan.

**Keywords:** arsenic, small ubiquitin-like modifier (SUMO), solubility, phase separation, promyelocytic leukemia-nuclear body (PML-NB), extranuclear PML body (EnPB)

## Abstract

Promyelocytic leukemia (PML) proteins are known to form phase-separated nuclear punctate structures called PML-nuclear bodies (PML-NBs). The integrity disruption of PML-NBs is linked with the pathogenesis of acute promyelocytic leukemia (APL), and trivalent arsenic (As^3+^) has been used for the clinical treatment of APL to restore normal PML-NBs. As^3+^ is considered to bind to cysteine residues and enhances modification of PML with small-ubiquitin-like protein (SUMO). We exposed U-2OS and CHO-K1 cells stably overexpressing PML-VI to As^3+^ and found that the solubility of PML decreased and SUMOylation of PML increased after 2 h-exposure to 3 μM As^3+^. Contrary to As^3+^-induced remarkable biochemical changes including the solubility change and SUMOylation of PML, microscopic observation of PML-NBs was not changed clearly after a short-term exposure to As^3+^. The number of PML-NBs decreased and extranuclear PML bodies (EnPBs), which are remniscences of PML-NBs after nuclear membrane breakdown at mitosis, increased after exposure to As^3+^ for 24 - 72 h. The amount of SUMOylated PML decreased after prolonged exposure to As^3+^ while the solubility of PML was kept low, suggesting that As^3+^ stabilized EnPB without SUMOylation. The effects of As^3+^ on EnPBs were clearly observed at as low as 0.3 μM As^3+^ which corresponds to inorganic arsenic level in drinking water worldwide.

## Introduction

It is known that trivalent arsenic (arsenite, As^3+^) is an effective chemotherapeutic agent for acute promyelocytic leukemia (APL) which is caused by the reciprocal t(15;17) translocation of promyelocytic leukemia (PML) and retinoic acid receptor-α (RARα) genes and the resultant expression of dominant negative PML-RARα fusion proteins (de The *et al*., 1991; Kakizuka *et al*., 1991). All PML isoforms (I to VII) are cysteine-rich proteins and consist of RING, two B-boxes, and coiled-coil domains called RBCC or tripartite motif, and PML isoforms (I to VI) have an NLS (Maroui *et al*., 2012; Nisole *et al*., 2013). As^3+^ is thought to bind to the cysteine residues changing conformation of PML and enhance degradation of PML-RARα by ubiquitin proteasome system (UPS) (Tatham *et al*., 2008).

Nuclear PML isoforms are scafold proteins of PML-NBs which are found as 10 to 30 dots of 0.2-1 μm in most mamallian cells (Brand *et al*., 2010). PML-NBs are reportedly formed by liquid-liquid phase separation (LLPS) (Corpet *et al*., 2020). SUMOylation at K65, K160, and K490 on PML and the SUMO-SIM interaction seem to play a role for the formation of PML-NBs (Li *et al*., 2017; Hoischen *et al*., 2018), but mutation at these lysine sites did not abolish PML-NB formation, suggesting that the SUMO-SIM interaction is not requisite for the dot-like assembly of PML proteins (Shen *et al*., 2006). The SUMO-SIM interaction does not work as a simple molecular glue because inhibition of SUMOylation by ML792 increased the size of PML-NBs (Hirano and Udagawa, 2022b).

The fragment of RING domain of PML forms supramolecular assemblies of ca. 50 nm in vitro which is similar to those formed by their full-length counterparts in cells. Mutation at the first zinc-binding site of RING fails to self-assemble PML *in vitro* and to function as translational repression in cells, suggesting the importance of RING domain for the formation of PML-NBs in cells (Kentsis *et al*., 2002). PML proteins prefer a dimer which extends long CC helices to the opposite direction like an octopus’s arms. Hydrophobic amino acids located in an α-helix franked with RING and b-BOX1 domains (F108, F109, L112, and L118) play a critical role in the PML monomer structure and PML-NB formation. The three hydrophobic pockets in the CC domain promote CC polymerization in an anti-parallel manner. (Tan *et al*., 2024).

Exposure to As^3+^ for 1-2 h dramatically reduces the solubility of PMLs in detergent-containing buffer solution and SUMOylates PML with both SUMO1 and SUMO2/3 in mammalian cells (Hirano *et al*., 2013; Hirano, 2020). The SUMOylated PMLs are subsequently ubiquitinated by SUMO-targeted ubiquitin ligases (STUbL) such as RNF4 (Lallemand-Breitenbach *et al*., 2008; Tatham *et al*., 2008) and TOPORS (Liu *et al*., 2024). However, the behavior of PML after a prolonged exposure to As^3+^, and the restoration of PML after removal of As^3+^ have not been well documented. One of the most explicit changes in PML-NBs caused by long exposure to As^3+^ is agglomeration of PML-NBs (Hirano and Udagawa, 2022a). The agglomeration of PML-NBs is also caused by antimony ion (Sb^3+^) which has a potency to insolubilize and SUMOylate PML proteins like As^3+^ (Muller *et al*., 1998; Hirano *et al*., 2015), suggesting that SUMOylation may play a role in the stability of PML-NBs.

PML-NBs agglomerate at mitosis and the agglomerated blobs, which are referred to as mitotic assemblies of PML proteins (MAPPs), complex with nucleoporins at the peri-nuclear region on the entry into G1 phase in daughter cells. The blobs at this stage is called cytoplasmic assemblies of PML and nucleoporins (CyPNs) (Jul-Larsen *et al*., 2009; Lang *et al*., 2017). Exposure to 0.5 μM arsenic trioxide (ATO) stabilized CyPNs with concomitant inhibition of PML-NB formation in HaCaT and NB4 daughter cells (Lang *et al*., 2012). However, biochemical analysis of PML after a long exposure to As^3+^ has not been reported except that CyPNs contain less SUMO molecules than PML-NBs. We, therefore, address the following three critical issues regarding phase separation of PML proteins in the present study. We refer to peri-nuclear PML agglomerates including CyPNs as extranuclear PML bodies (EnPBs) because the association of extranuclear PML with nuclear pores was not confirmed in the present study.

1. Regulation of PML-NBs by SUMOylation.
2. Effects of As^3+^ on the assembly of PML-NBs.
3. Effects of As^3+^ on the stability of EnPBs.

## Materials and Methods

### Chemicals and antibodies

Sodium *m*-arsenite and other common chemicals were obtained from Sigma-Aldrich (St Louis, MO) or Fujifilm-Wako (Osaka, Japan). BCA protein assay kit was obtained from Thermo Fisher (Waltham, MA). ML792 (SUMO E1 inhibitor) and TAK294 (ubiquitin E1 inhibitor) were obtained from MedKoo Bioscience (Morrisville, NC) and Selleckchem (Houston, TX), respectively. Details for antibodies used for immunoblotting and immunostaining, and plasmids used for transfection were provided as supplementary information (Source table). Alexa Fluor™ 594-labeled anti-SUMO2/3 antibody was used for immunostaining. The conjugated antibody was prepared using an antibody labeling kit with Alexa Fluor (Thermo Fisher).

### Cells

U-2OS cells (HTB-96) were obtained from ATCC and cultured in McCoy’s 5A medium containing 10% heat-inactivated fetal bovine serum (FBS). The cells were transfected with *PML-VI* plasmid (RC220236, transcript variant 5, Origene, Rockville, MD), and cells stably expressing PML-VI were selected by G418 (*PML(VI)*-U2OS cells). *PML*-null U-2OS cells were prepared using a double-nickase vector (Santa Cruz, Dallas, TX) and the following selection with puromycin (*PML^-/-^*U-2OS). CHO-K1 (RCB0285) and HEK293 cells (RCB1637) were obtained from RIKEN (Ibaraki, Japan), and were cultured in complete Ham’s F12 medium and DMEM, respectively. CHO-K1 cells stably expressing GFP-tagged human PML-VI (Hirano *et al*., 2015) were further transfected with mChery-tagged *SUMO2* plasmids (wild, and K11R and ΔGG mutants; VectorBuilder Inc., Yokohama, Japan), and stable cell lines were selected by puromycin (CHOgPrS cells).

### Western blotting

The cells were washed with HBSS and lysed with ice-cold RIPA buffer containing protease (Santa Cruz) and phosphatase inhibitor cocktails (Calbiochem, San Diego, CA) on ice for 10 min. The cell lysate was centrifuged at 9,000 *g* for 5 min at 4 °C. The supernatant was collected and labeled as the RIPA-soluble fraction (Sol). The pellet was rinsed with PBS and treated with Benzonase^®^ nuclease (240 U/mL in Tris-HCl buffer, Santa Cruz) solution of the same volume as the RIPA buffer at 25 °C for 2 h with intermittent mixing (Thermomixer comfort, Eppendorf, Wesseling-Berzdorf, Germany). The digested sample was labeled as the RIPA-insoluble fraction (Ins). Protein concentrations of the RIPA soluble fractions were adjusted at 3.5 mg/mL. Aliquots of the RIPA-soluble and the RIPA-insoluble fractions were mixed with LDS sample buffer (1x TBS, 10% glycerol, 0.015% EDTA, 50 mM DTT, and 2% LDS) and heated at 95 °C for 5 min. The samples were resolved by LDS (SDS)-PAGE (4-12%) with MES buffer and electroblotted onto a PVDF membrane. The membranes were first stained with Ponseau S (Santa Cruz) and blocked with PVDF Blocking Reagent (TOYOBO, Osaka, Japan) after removing Ponseau S in 0.01 N NaOH solution. The antibodies were diluted using Can-Get-Signal^TM^ solutions (TOYOBO, Osaka, Japan). The chemiluminescent signals were captured with a CCD camera (ImageQuant 800, GE Healthcare, Uppsala, Sweden) after soaking the membrane in ECL (Prime, GE Healthcare, Buckinghamshire, UK). The membrane was stripped in warmed (56 °C) Tris-HCl buffer (62.5 mM, pH 6.7) containing 100 mM 2-mercaptoethanol and 2% SDS with vigorous shaking for 3 min and blocked again for the second probing. Finally, the membrane was reprobed again for the detection of tubulin and histone H3 to assess the loaded protein levels in the RIPA-soluble and the RIPA-insoluble fractions, respectively. The densitometric analyses of the membranes were performed using ImageJ software (https://imagej.nih.gov/ij/).

### Live cell imaging

CHOgPrS cells were pre-cultured in an 8-well cover-glass chamber (Iwaki, Tokyo). After exposure to As^3+^ or the other chemicals, fluorescent images of the cells was captured by microscopy (BZ-X800, Keyence, Osaka). NucBlue™ Live ReadyProbes™ Reagent (Hoechst 33342, Thermo Fisher) was applied to visualize each nucleus.

### Immunostaining

U-2OS cells were grown in an 8-well chamber slide (Millicell EZ slide, Merck Millipore, Burlington, MA) to early confluency. The cells were washed with warmed (37 °C) HBSS and fixed with 3.7% formaldehyde solution for 10 min, permeabilized with 0.1% Triton X-100 for 10 min, and treated with Image-iT™ FX Signal Enhancer (Thermo Fisher) for another 10 min. The cells were immunostained with rabbit polyclonal anti-PML antibody followed by Alexa Fluor 488-labeled goat anti-rabbit IgG, then further stained with Alexa Fluor 594-labeled anti-SUMO-2/3. Can-Get-Signal^TM^ Immunostain solutions (TOYOBO) were used to dilute the antibodies. The cell nuclei were counterstained with DAPI (NucBlue™ Fixed Cell ReadyProbes™ Reagent, Thermo Fisher). Digital fluorescence images were captured using a fluorescence microscope (Eclipse 80i, Nikon, Tokyo), and the images were assembled using Photoshop® software (Adobe, San Jose, CA).

### Statistical analyses

Data were presented as means ± SD for the numbers of PML-NBs and EnPBs. Densitometric data were presented as means ± SEM. Statistical analyses were performed by one-way ANOVA followed by Tukey’s *post-hoc* test. A probability value less than 0.05 was accepted as indicative of statistical significance.

## Results

### Formation of PML-NBs and EnPBs in U-2OS cells

U-2OS cells were exposed to 3 μM As^3+^ for 2 and 72 h, and the cells were immunostained with anti-PML and anti-SUMO2/3 antibodies. PML-NBs, which are characterized by co-localization of PML and SUMO2/3 in nuclei, were visible in untreated cells. PML-NBs were more clearly observed after 2-h exposure to As^3+^. However, PML-NBs were scarcely observed after exposure to As^3+^ for 72 h and EnPBs increased reciprocally (Fig. 1(A)). Western blot analysis indicated that at least three PML isoforms were detected in the untreated cells and PML proteins became insoluble in the RIPA buffer and SUMOylated by 2 h-exposure to 3 μM As^3+^. These findings were not clearly observed by 2 h-exposure to 0.3 μM As^3+^. PML proteins in the soluble fraction decreased after 72 h-exposure to either 0.3 or 3 μM As^3+^ (Fig. 1(B)), and were restored in the RIPA-soluble fraction 24 h after removing As^3+^ from culture medium (Supplementary Fig. 1(A)). These As^3+^-induced changes in endogenous PML proteins were also observed in HEK293 cells (Supplementary Fig. 1(B)).

**Fig. 1.**
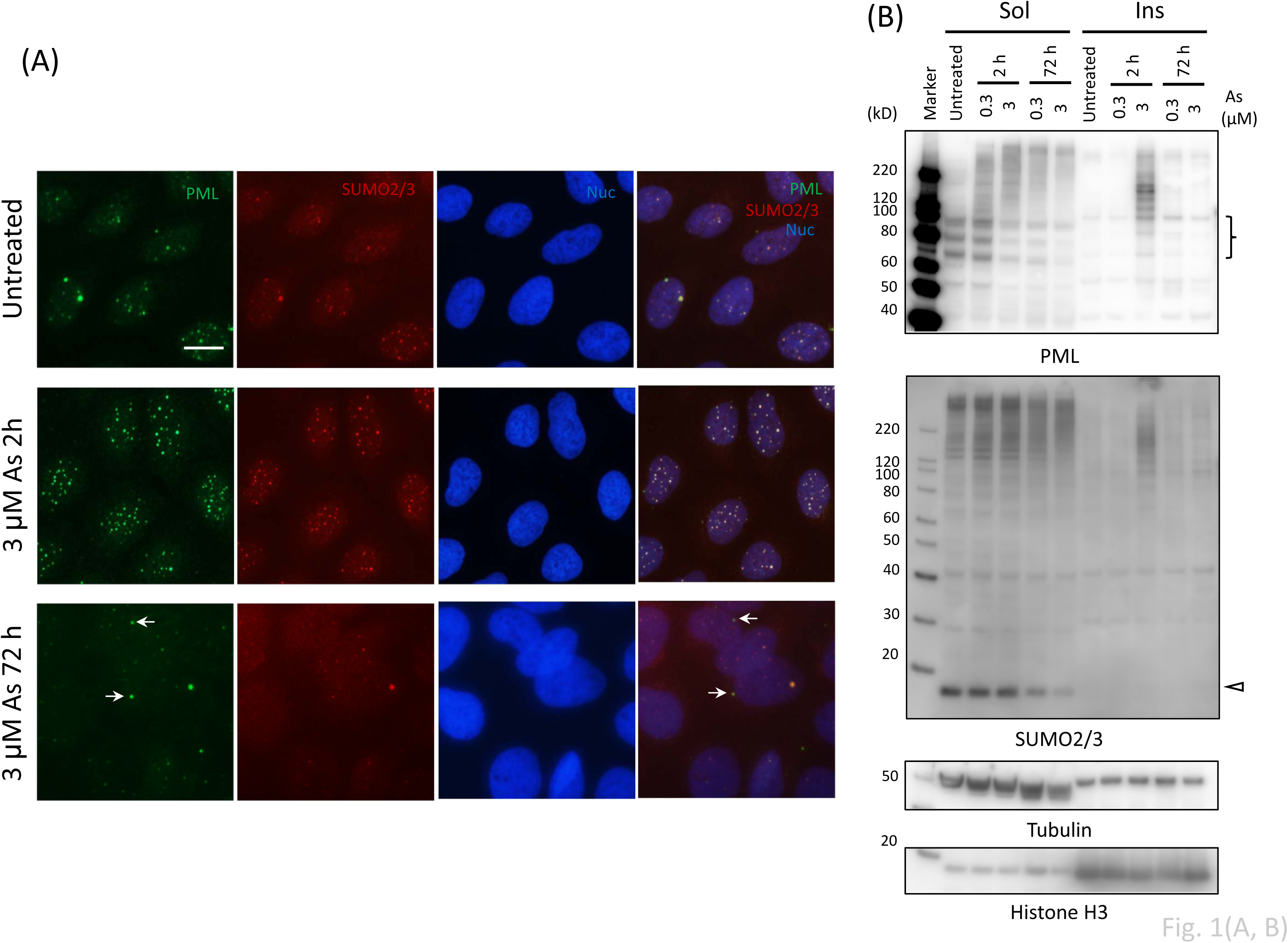
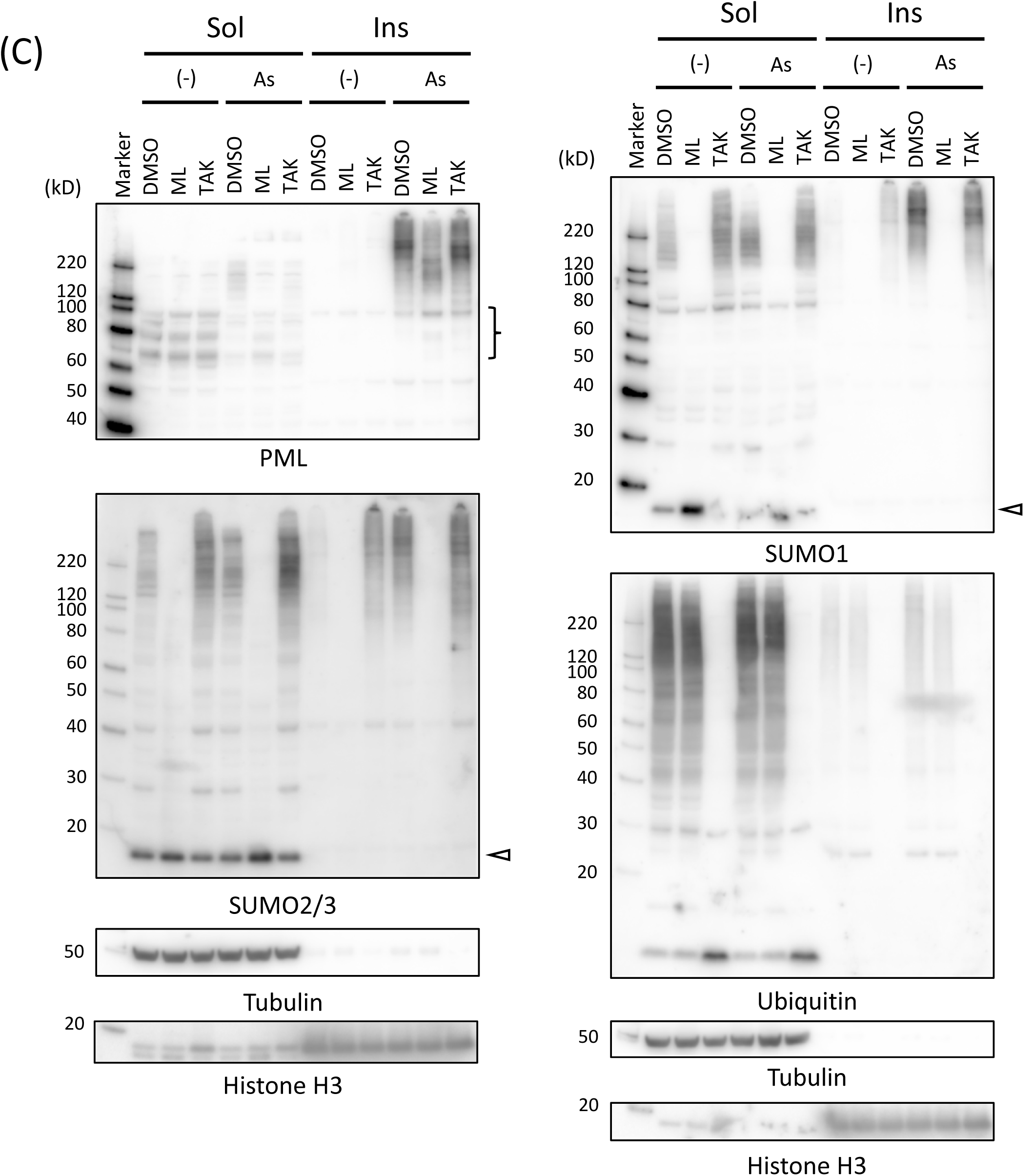
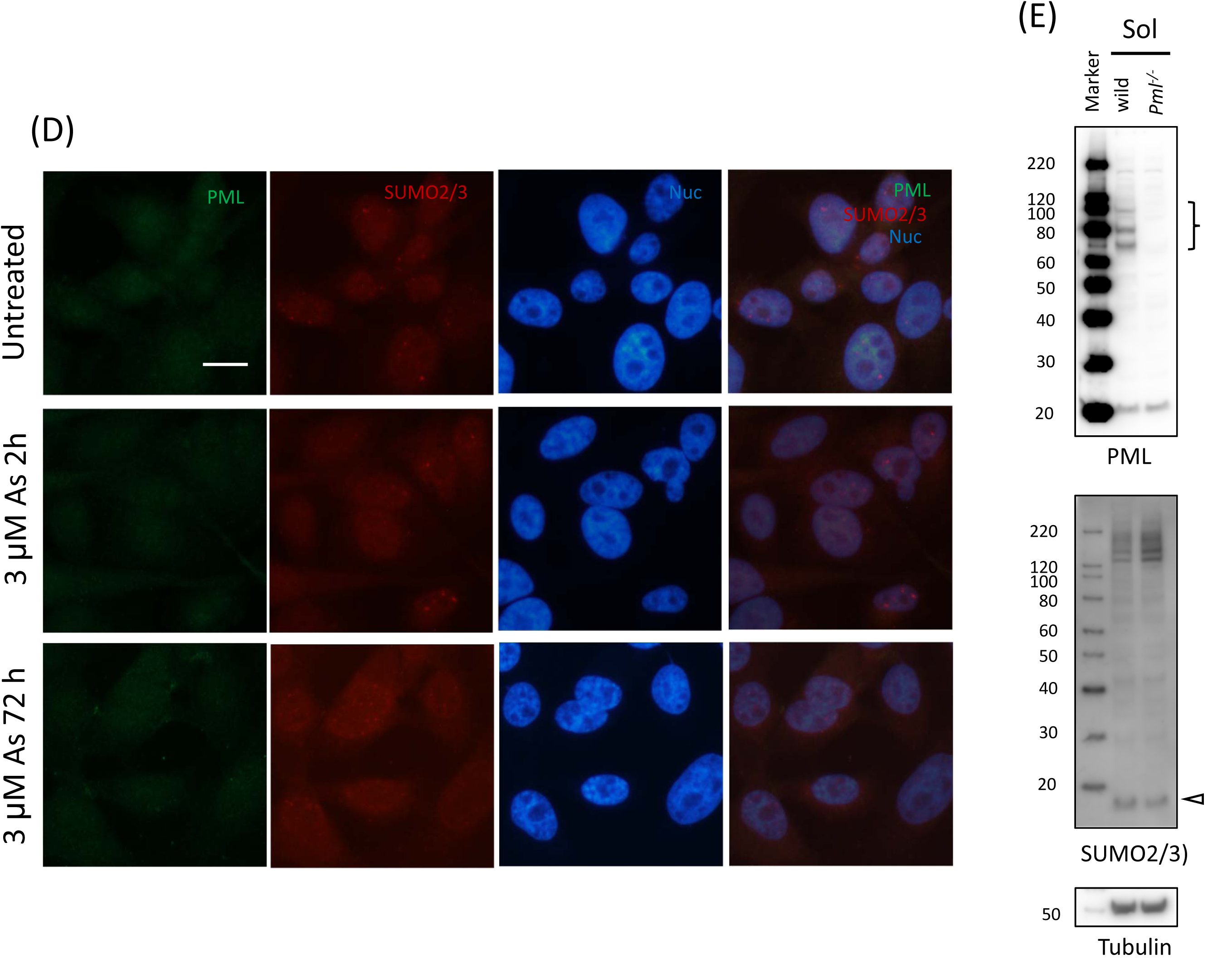
Effects of As^3+^ on intranuclear and extranuclear PML in U-2OS cells. (A) Immunofluorescence micrographs of U-2OS cells. The cells were exposed to 3 μM As^3+^ for 2 or 72 h, or left untreated. After immunostaining for PML and SUMO2/3 antibodies, the cells were counterstained with DAPI (Nuc). Scale bar = 20 μm. An arrow indicates an EnPB. (B) Western blot analyses for PML and SUMO2/3 in U-2OS cells. The cells were treated with either 0.3 or 3 μM As^3+^ for 2 or 72 h, or left untreated, and then lyzed with ice-cold RIPA buffer. The lysate was separated into soluble and insoluble fractions by centrifugation. (C) Western blot analyses for PML, SUMO2/3, SUMO1, and ubiquitin in U-2OS cells. The cells were pretreated with 20 μM ML792 (SUMO E1 inhibitor) or 10 μM TAK294 (ubiquitin E1 inhibitor) for 3 h, and were exposed to 3 μM As^3+^ for 2 h in the presence of the inhibitor. (D) Immunofluorescence micrographs of *PML^-/-^* U-2OS cells. The cells were exposed to 3 μM As^3+^ for 2 or 72 h, or left untreated. Scale bar = 20 μm. (E) Western blot analyses for PML and SUMO2/3 in the soluble fractions of wild U-2OS and *PML^-/-^*U-2OS cells. Note that SUMOylated proteins were detected in low electromobility region in both cell types. An open arrowhead indicates SUMO monomers. A right curly bracket indicates PML isoforms.

Pre-exposure to ML792, a SUMO E1 activating inhibitor, significantly reduced SUMOylation of proteins with SUMO1 and SUMO2/3 in both RIPA-soluble and RIPA-insoluble fractions irrespective of As^3+^ exposure. TAK243, a ubiquitin E1 activating inhibitor, diminished ubiquitinated proteins and did not affect the soluble-to-insoluble shift of PML proteins, suggesting that ubiquitination did not affect the As^3+^-induced biochemical changes of PML proteins, if any (Fig. 1(C)). Neither PML-NBs nor EnPBs were observed as clear dots in *PML*-null cells (*PML^-/-^*U-2OS). SUMO2/3, which expressed equally in both types of U-2OS cells, distributed diffusely in the cells (Fig. 1(D) and (E)). The SUMOylated PML proteins in the RIPA-insoluble fraction gradually disappeared in U-2OS and HEK293 cells following continuous exposure to 3 μM As^3+^ for 24 h (Supplementary Fig. 1(A) and (B)). These results indicate that PML proteins were hardly recycled as the scaffold of PML-NBs in the presence of As^3+^.

### Formation of PML-NBs and EnPAs in U-2OS cells stably overexpressing PML-VI (*PML(VI)* U-2OS cells)

PML-VI-overexpressing cells were used to visualize time-course changes in biogenesis of PML-NBs and EnPBs clearly. Both PML-NBs and EnPBs were observed as large blobs in *PML(VI)* U-2OS cells. The number of PML-NBs decreased and that of EnPBs increased reciprocally 72 h after exposure to As^3+^, but in contrast to U-2OS cells they were still visible clearly in *PML(VI)* U-2OS cells (Fig. 2(A) and (B)). PML-VI in the RIPA-soluble fraction decreased 2 h after exposure to 3 μM As^3+^, and PML-VI and SUMOylated PML-VI bands were detected in the RIPA-insoluble fraction. Although the intensity of PML-VI band was not changed up to 72 h, SUMOylated PML-VI decreased with time (Fig. 2(C)), suggesting that SUMOylation does not play an important role for As^3+^-induced EnPBs in *PML(VI)* U-2OS cells. Quantitative analysis of densitometric data indicated that SUMO2/3 and SUMO1 monomers, which were detected in the RIPA-soluble fraction, decreased 2 h after exposure to 3 μM As^3+^ along with the soluble-to-insoluble shift of PML-VI (Supplementary Fig. 2).

**Fig. 2.**
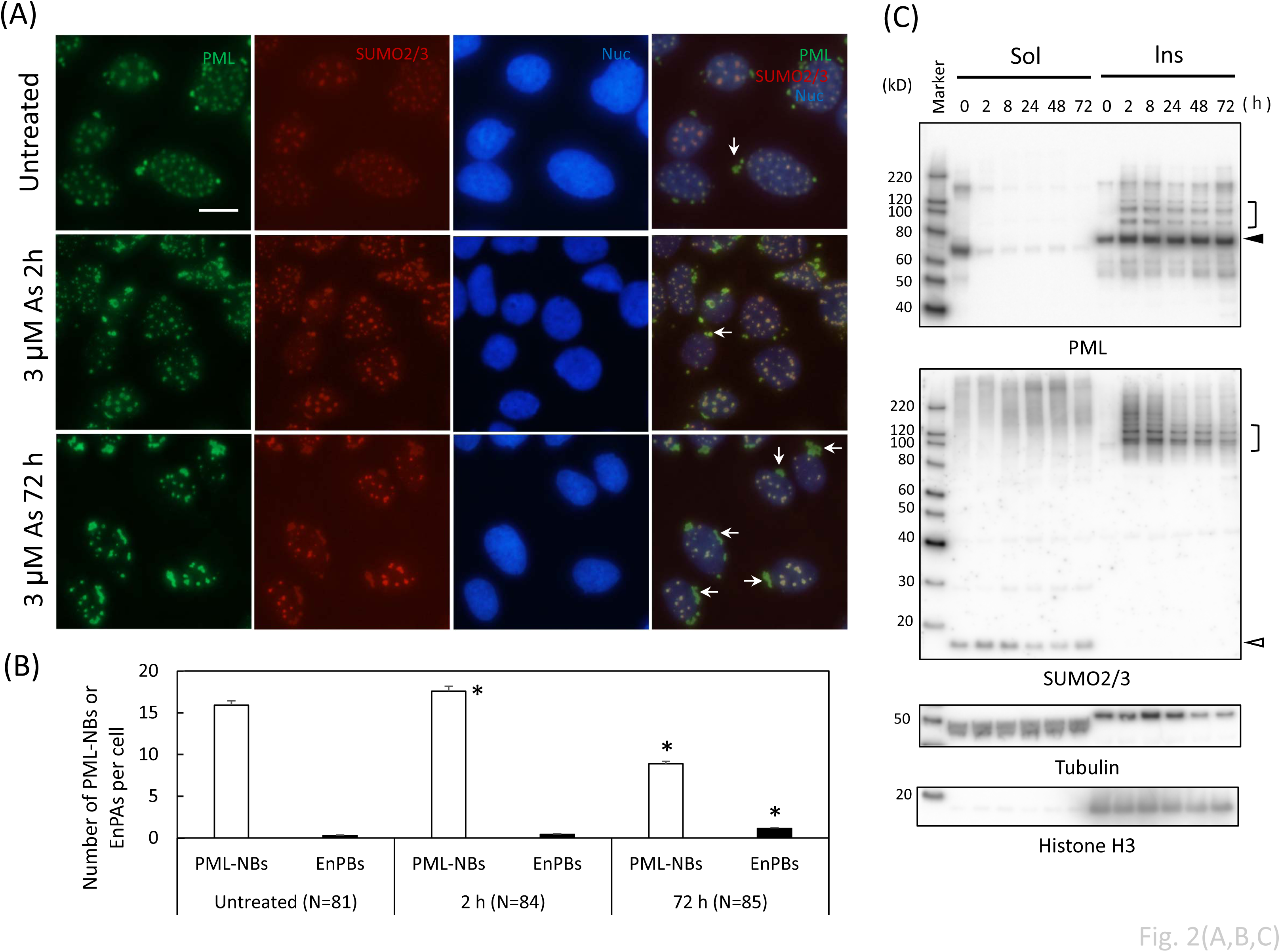
Effects of As^3+^ on intranuclear and extranuclear PML in *PML(VI)* U-2OS cells. (A) Immunofluorescent micrographs of *PML(VI)* U-2OS cells. The cells were exposed to 3 μM As^3+^ for 2 or 72 h, or left untreated. Scale bar = 20 μm. An arrow indicates an EnPB. (B) Numbers of PML-NBs and EnPBs in untreated and As^3+^-exposed *PML(VI)* U-2OS cells. An agglomeration of three or more than three PML blobs in the perinuclear region was counted as an EnPB. More than 80 cells were observed for counting PML-NBs and EnPBs. Data are presented as mean ± SD. *, Significantly different from the untreated cells. (C) Western blot analysis for PML and SUMO2/3 following exposure to 3 μM As^3+^ for 0, 2, 8, 24, 48, or 72 h in *PML(VI)* U-2OS cells. Closed and open arrowheads indicate PML-VI and SUMO2/3 monomers, respectively. A right square bracket indicates SUMOylated PML-VI.

### PML-NBs and EnPBs in As^3+^-exposed CHOgPrS cells

GFP-tagged PML-VI proteins were present as PML-NBs in nuclei in untreated CHOgPrSwild and CHOgPrSΔGG cells. Wild-type SUMO2 molecules were co-localized with PML-NBs as dense dots, while ΔGG-mutant SUMO2 was present diffusely in the cells. The distribution of GFP-tagged PML and mCherry-tagged SUMO2 were not changed largely after 2-h exposure to 3 μM As^3+^ except that wild-type SUMO2 was more limited to PML-NBs (Fig. 3 (A and B)). Exposure to 3 μM As^3+^ for 2 h changed GFP-tagged PML-VI from the RIPA-soluble to the RIPA-insoluble form, and GFP-tagged PML-VI was SUMOylated with endogenous SUMO2/3 in CHOgPrSΔGG cells and mostly with mCherry-tagged SUMO2 in CHOgPrSwild and CHOgPrS(K11R) cells (Supplementary Fig. 3 (A)). However, the number of PML-NBs decreased and most GFP-tagged PML-VI were observed as EnPBs at peri-nuclear regions after 24 h in both CHOgPrSwild and CHOgPrSΔGG cells. Notably, EnPBs contained much less SUMO2 molecules than PML-NBs (Fig. 3(C) and (D)). The formation of EnPBs was not observed by exposure to Cd^2+^ or Cu^2+^ (Fig. 3(E) and Supplementary Fig. 4). Exposure to 3 μM Cd^2+^ reduced the number of PML-NBs with shrinking of cells, suggesting that the decreased number of PML-NBs might be due to cytotoxic effects of Cd^2+^. EnPBs were also observed by exposure to 3 μM Sb^3+^ (Supplementary Fig. 4).

**Fig. 3.**
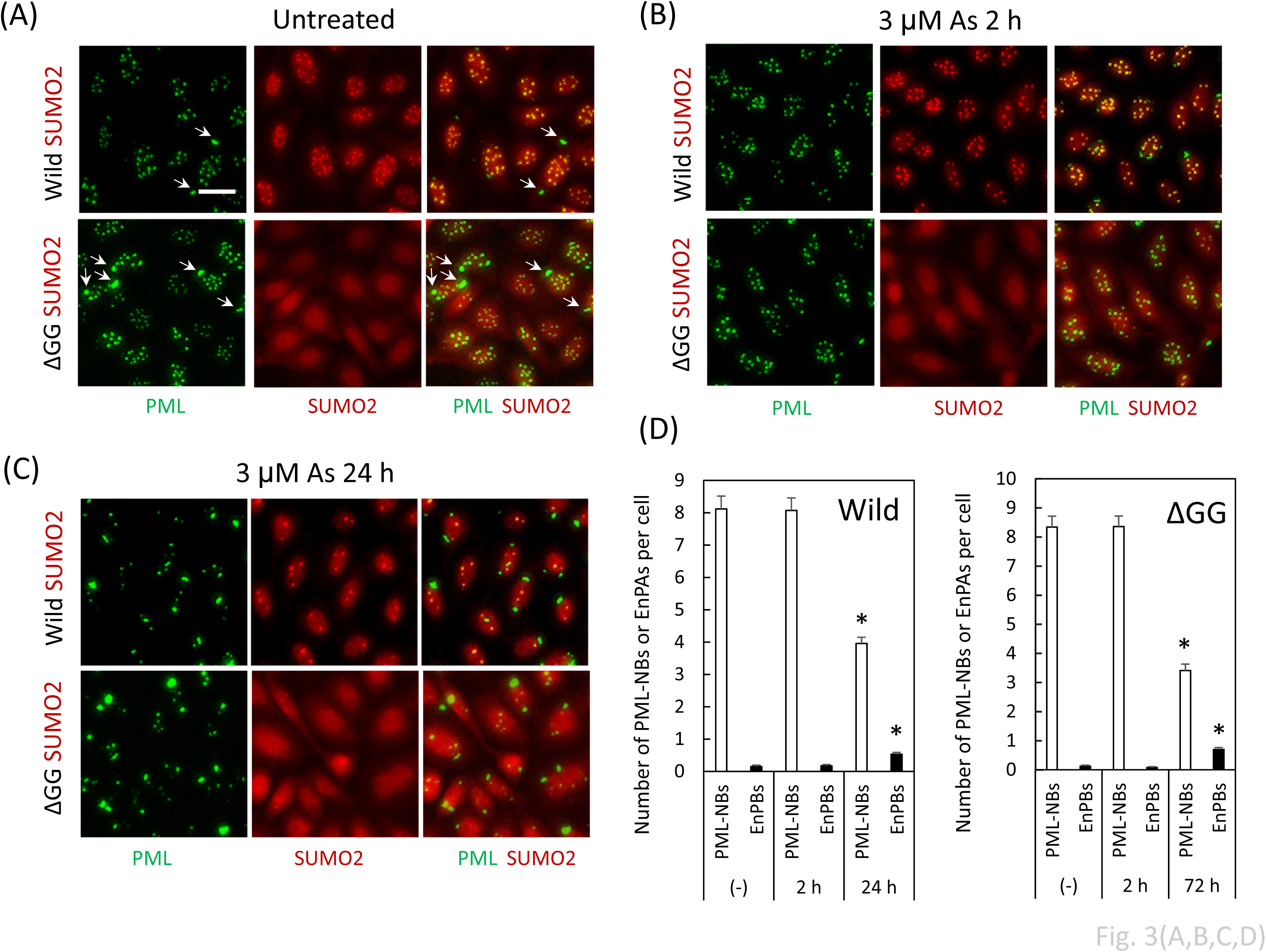
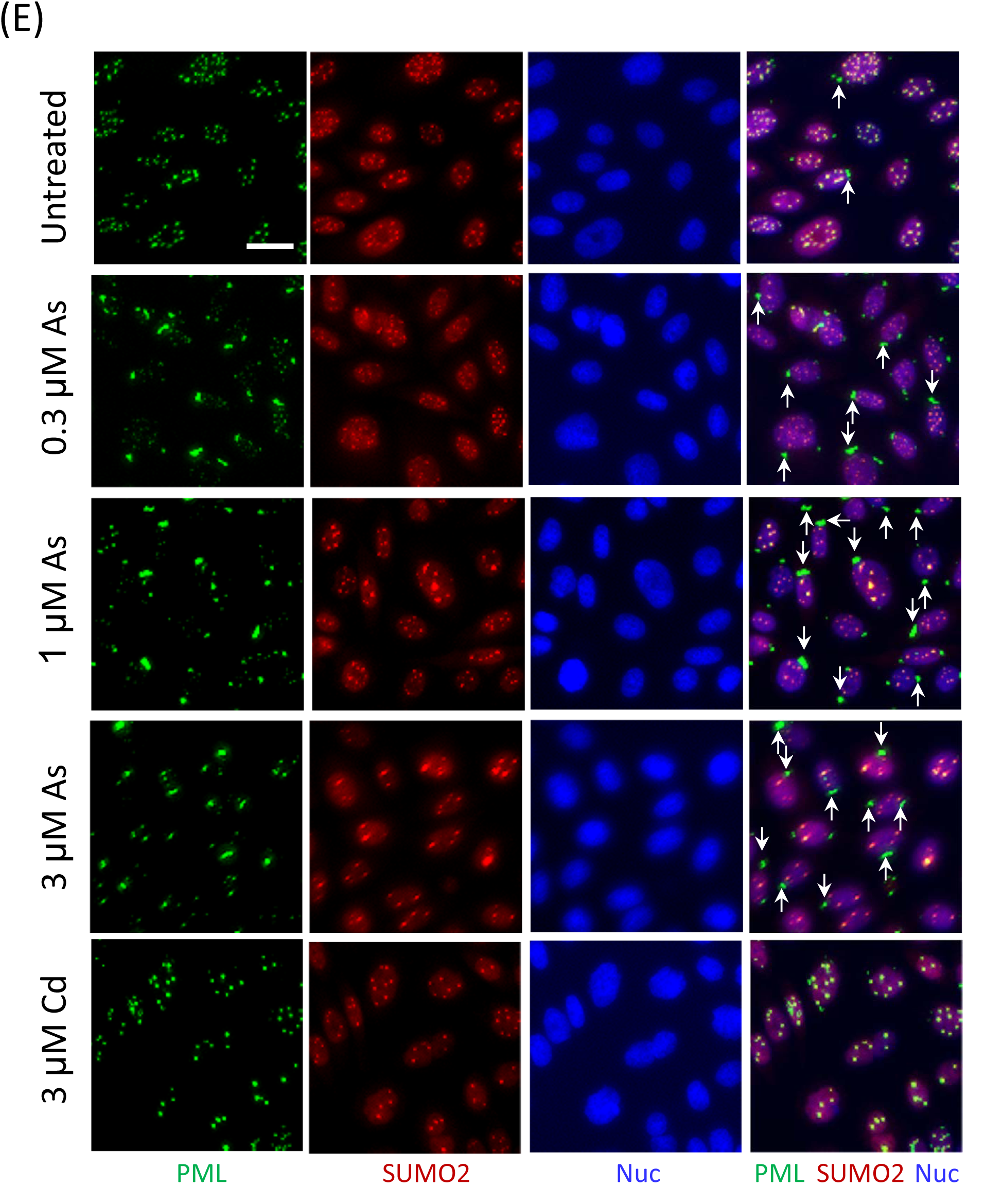
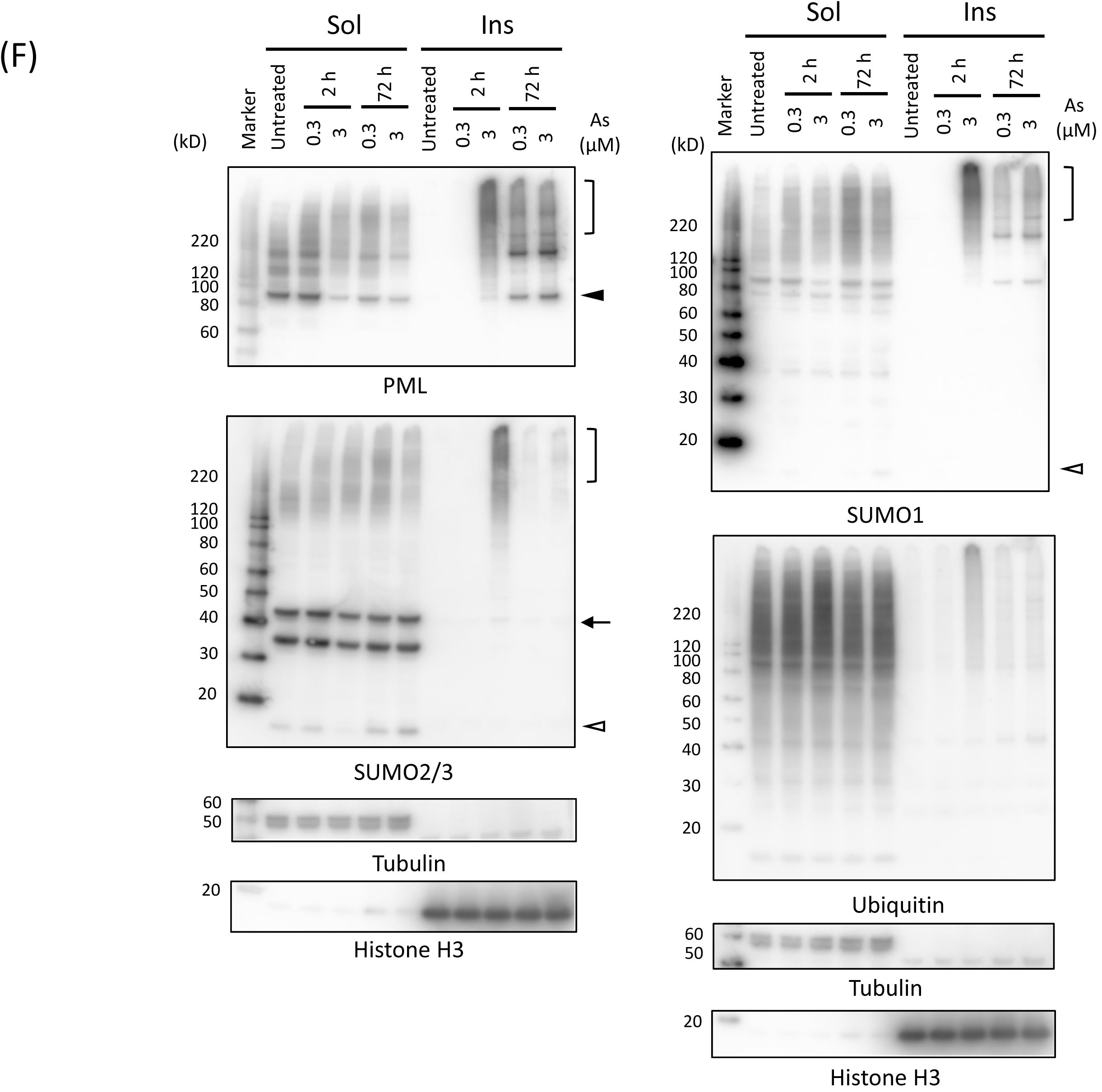
Changes in PML-NBs and extranuclear PML bodies (EnPBs) in CHO-K1 cells stably expressing GFP-PML-VI and wild-type (CHOgPrSWild) or C-terminal diglycine-deleted mCherry SUMO2 (CHOgPrSΔGG) cells. (A-C) Fluorescent micrographs of CHOgPrSWild cells. The cells were exposed to 3 μM As^3+^ for 2 h (B) or 24 h (C), or left untreated (A). Scale bar = 20 μm. An arrow indicates an EnPB. (D) Numbers of PML-NBs and EnPBs in untreated and As^3+^-exposed CHOgPrSWild and CHOgPrSΔGG cells. More than 90 cells were observed for counting PML-NBs and EnPBs. Data are presented as mean ± SD. *, Significantly different from the untreated CHOgPrSWild or CHOgPrSΔGG cells. (E) Fluorescent micrographs of CHOgPrSWild cells. The cells were exposed to 0.3, 1, or 3 μM As^3+^ or 3 μM Cd^2+^ for 24 h or left untreated. Scale bar = 20 μm. An arrow indicates an EnPB. (F) Western blot analyses for SUMO2/3, PML, SUMO1, and ubiquitin in the RIPA-soluble and -insoluble fractions of CHOgPrSWild cells. The cells were exposed to 0.3 or 3 μM As^3+^ for 2 or 72 h. An open arrowhead indicates SUMO monomer. A closed arrowhead indicates GFP-tagged PML-VI. An arrow indicates an mCherry-tagged SUMO2. A right square bracket indicates SUMOylated GFP-tagged PML-VI.

Western blot analysis showed that 2 h-exposure to 3 μM As^3+^ turned the soluble GFP-tagged PML-VI into the insoluble form and SUMOylated GFP-tagged PML-VI with both SUMO2/3 and SUMO1. The insoluble form of GFP-tagged PML-VI was only slightly ubiquitinated. Exposure to 0.3 μM As^3+^ for 2 h did not cause these changes to GFP-tagged PML-VI. However, after exposure to either 0.3 μM or 3 μM As^3+^ for 3 days, GFP-tagged PML-VI was detected as low mobility products with an intense band migrating at around 170 kD together with an unmodified GFP-tagged PML-VI at 85 kD (Fig. 3(F)). It is reasonable to suppose that the low mobility products detected by western blotting correspond to major components of EnPBs in immunostaining.

### Effects of ML792 on SUMOylation of GFP-tagged PML-VI and the stability of EnPBs in As^3+^-exposed CHOgPrSwild cells

mCherry-tagged SUMO2 molecules associated with PML-NBs in untreated CHOgPrSwild cells. However, the association with PML-NBs was lost by ML792. Exposure to 3 μM As^3+^ did not recruit mCherry-tagged SUMO2 into PML-NBs in ML792-pretreated cells. However, pretreatment with 3 μM As^3+^ inhibited the effect of ML792 on dissociation of mCherry-tagged SUMO2 from PML-NBs. Agglomerated PML-NBs were observed following prolonged exposure to ML792 in As^3+^-pretreated cells (Fig. 4(A)). SUMOylated GFP-tagged PML-VI was present in the RIPA-soluble fraction as well as in the RIPA-insoluble fraction. Exposure to 3 μM As^3+^ for 2 h converted the soluble GFP-tagged PML-VI into the insoluble form. Pretreatment with ML792 inhibited SUMOylation of GFP-tagged PML-VI, however, the solubility shift of GFP-tagged PML-VI still occurred by exposure to 3 μM As^3+^. Comparison in SUMOylated GFP-tagged PML-VI between CHOgPrSwild and CHOgPrSΔGG cells indicated that endogenous and mCherry-tagged SUMO2 molecules competitively SUMOylated GFP-tagged PML-VI (Supplementary Fig. 3(B)). ML792 did not effectively reduce SUMOylation of GFP-tagged PML-VI in As^3+^-pretreated CHOgPrSwild cells, suggesting that SUMOylated GFP-tagged PML-VI was stabilized by As^3+^ and became resistant against deSUMOlation (Fig. 4(B)). These results are consistent with the microscopic observation that mCherry-tagged SUMO2 in PML-NBs were not affected by ML792 in As^3+^-pretreated CHOgPrSwild cells (Fig. 4(A)).

**Fig. 4.**
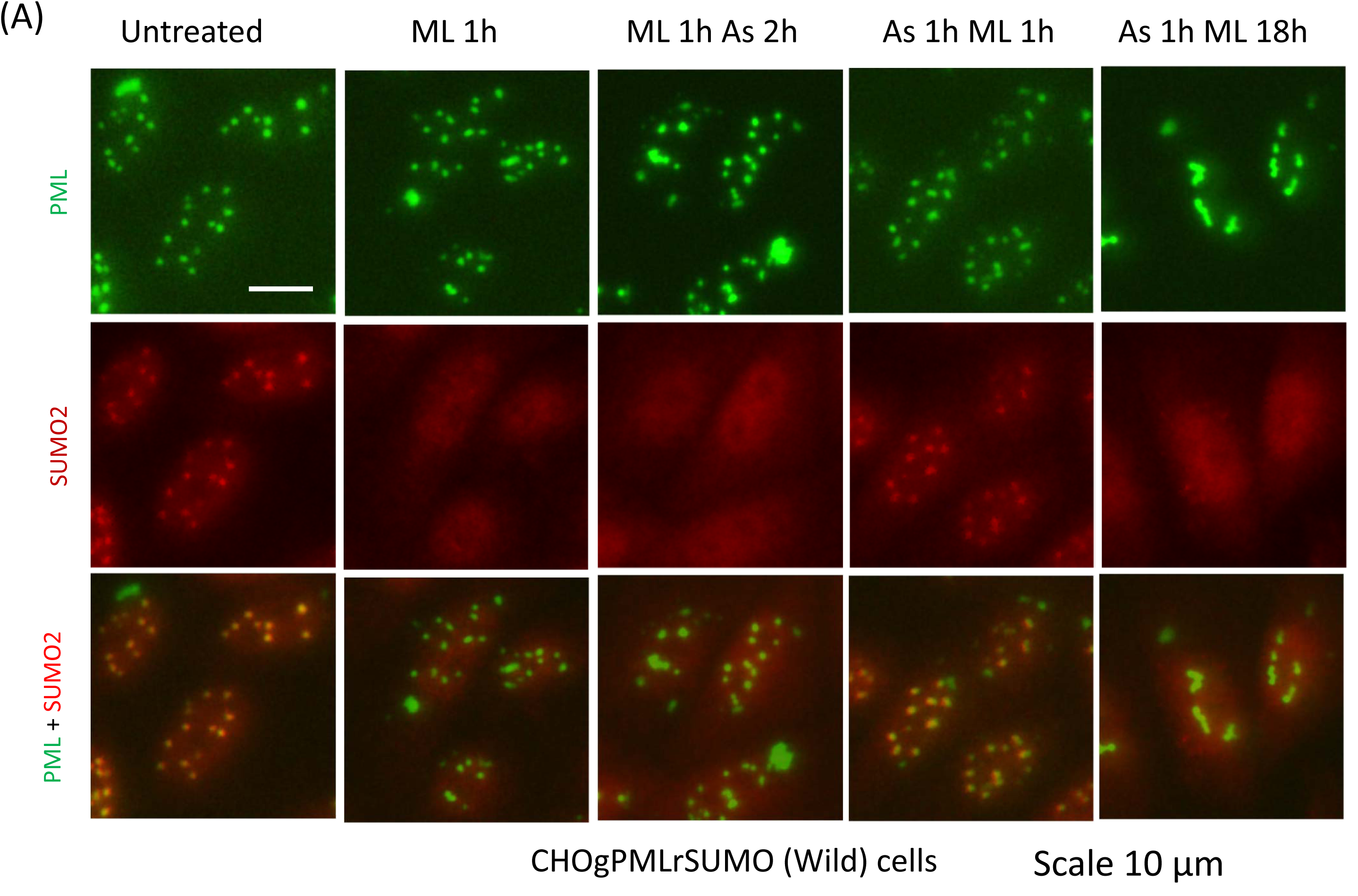
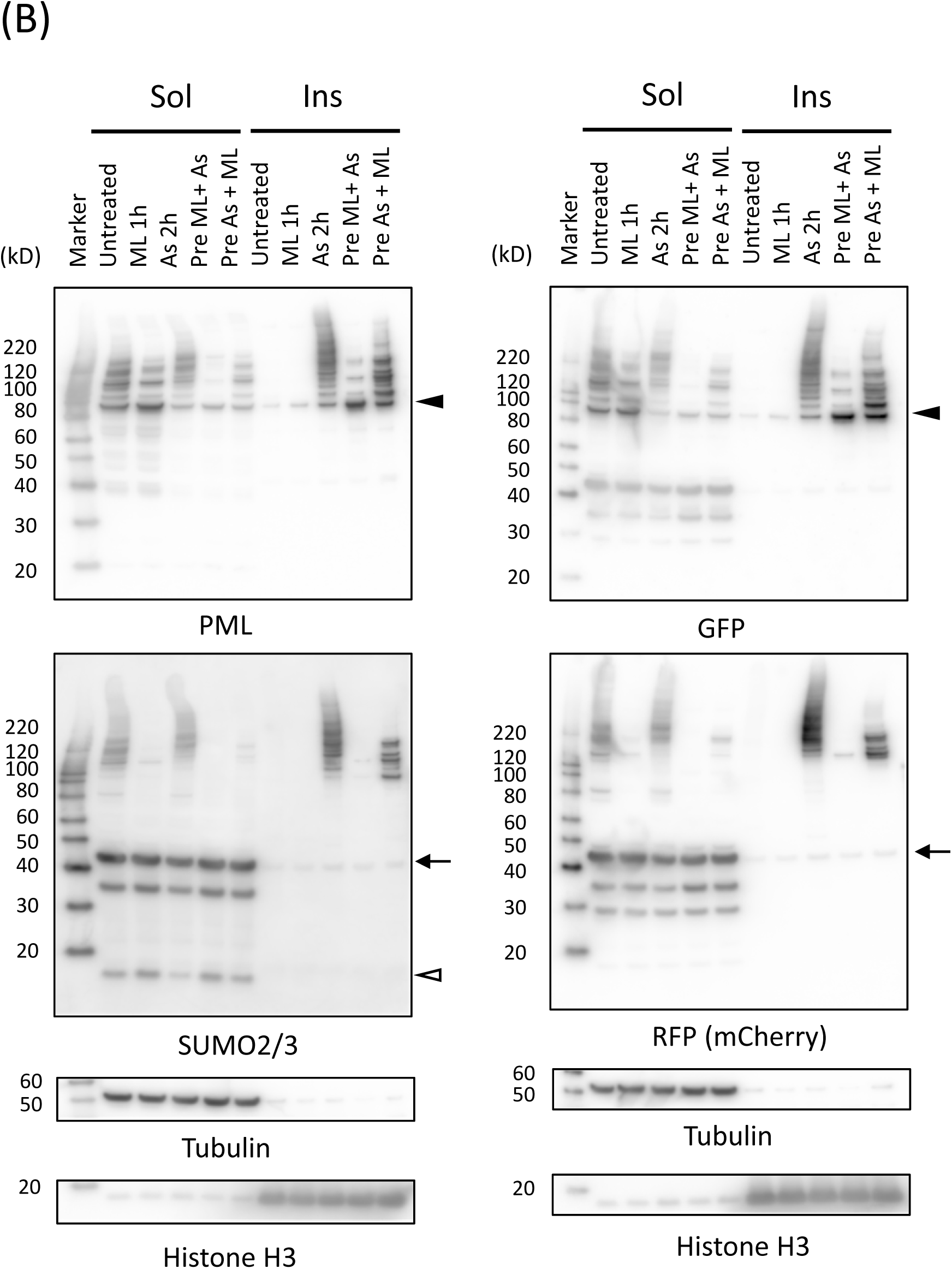
Effects of pre- and post-exposure of ML792 (SUMO E1 inhibitor) on As^3+^-induced changes in PML proteins and PML-NBs in CHOgPrSWild cells. (A) Fluorescent micrographs of CHOgPrSWild cells. The cells were pretreated with 20 μM ML792 for 1 h and exposed to 3 μM As^3+^ for 2 h, or pretreated with 3 μM As^3+^ for 1 h and exposed to ML792 for 1 or 18 h. Scale bar = 10 μm. (B) Western blot analyses for PML, SUMO2/3, GFP, and RFP (mCherry) in the RIPA-soluble and -insoluble fractions of CHOgPrSWild cells. The cells were pretreated with 3 μM As^3+^ for 2 h or 20 μM ML792 for 1 h, and then exposed to ML792 for 1 h or As^3+^ for 2 h, respectively. Closed and open arrowheads indicate GFP-tagged PML-VI and SUMO2/3 monomers, respectively. An arrow indicates an mCherry-tagged SUMO2.

The effect of As^3+^ on SUMOylation of PML was reversible, and SUMO2/3 was restored in PML-NBs after removal of As^3+^ from culture medium in CHOgPrSwild cells. Even though PML-NBs were restored in the nuclei after removal of As^3+^, EnPBs remained at perinuclear regions (Fig. 5(A)). Fig. 5(B) indicates that the intensity of the anti-PML antibody-reacting protein band at 170 kD was not changed even after 24 h after removal of As^3+^, suggesting that GFP-tagged PML-VI was converted to stable and yet unkown forms in As^3+^-induced EnPBs by As^3+^.

**Fig. 5.**
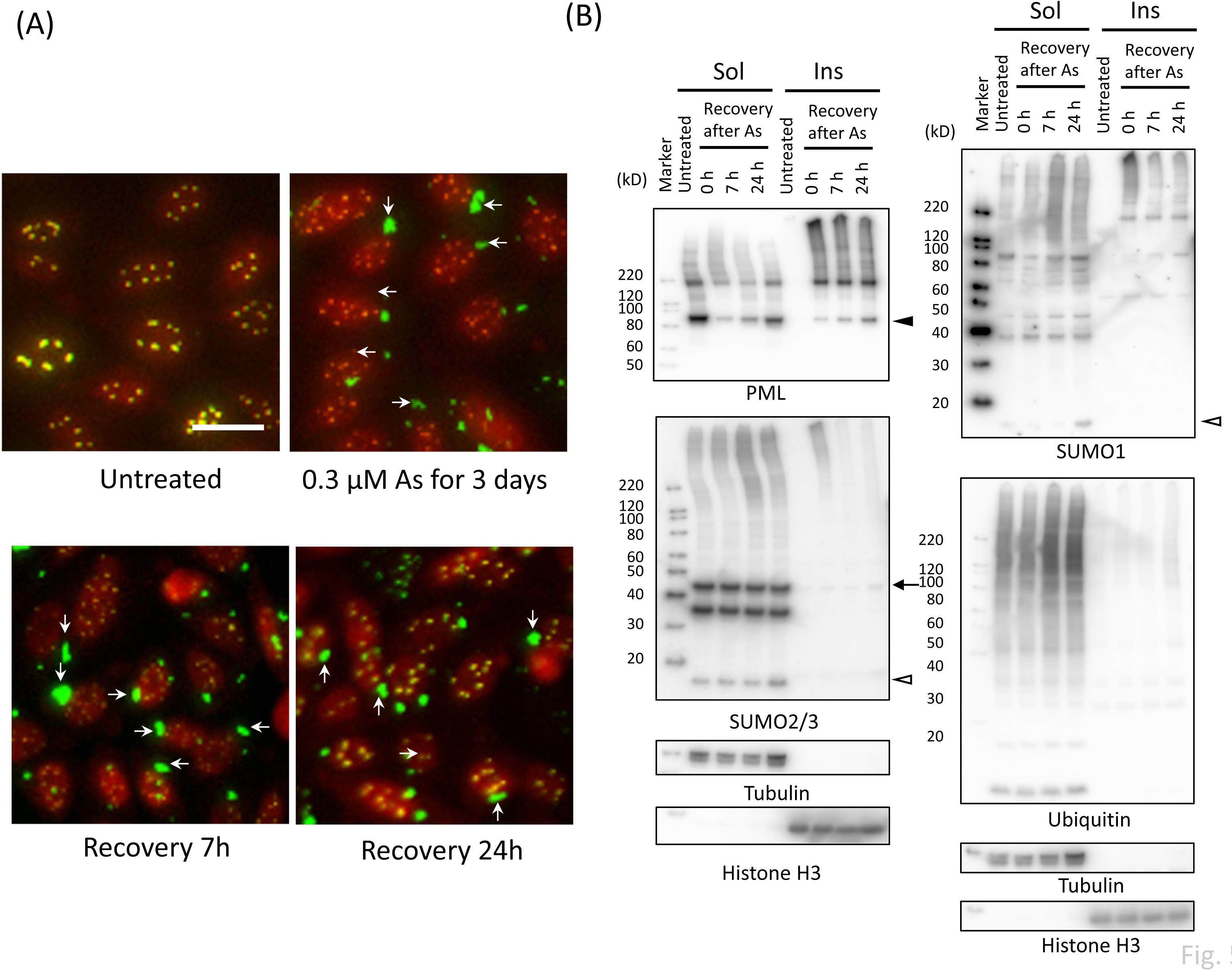
Recovery of PML-NBs and PML proteins after removal of As^3+^ from culture medium in CHOgPrSWild cells. (A) Fluorescent micrographs of CHOgPrSWild cells. The cells were exposed to 0.3 µM As^3+^ for 3 days, and further cultured in As^3+^-free fresh complete culture medium for 7 or 24 h. (B) Western blot analyses for PML, SUMO2/3, SUMO1, and ubiquitin in the RIPA-soluble and -insoluble fractions of CHOgPrSWild cells in the recovery from exposure to 0.3 µM As^3+^ for 3 days. See also the legend to (A).

## Discussion

There seem to be two distinctive PML forms in mammalian cells from the biochemical point of view, the cold RIPA-soluble and -insoluble forms. The soluble-to-insoluble change occurs shortly after exposure to As^3+^, and this process is followed by SUMOylation of PML proteins. However, these remarkable biochemical changes caused by As^3+^ are not reflected precisely in microscopic images of PML because PML and SUMO molecules were co-localized in PML-NBs of untreated cells, and the fluorescent images of PML-NBs appeared to be unchanged after 2 h-exposure to As^3+^ except that SUMO molecules looked more limited to PML-NBs (Figs. 1, 2 and 3). We hypothesized that SUMOylation and deSUMOylation on PML occur continuously and rapidly in untreated cells, and As^3+^ shifts this equilibrium toward SUMOylation. We found, for the first time, that biochemical characteristics of EnPBs are different from those of PML-NBs as shown in Figs. 3(F) and 5(B). Effects of As^3+^ on PML-NBs and EnPBs are schematically depicted in Fig. 6. Cellular uptake of As^3+^ through transporters (Liu *et al*., 2002) and intracellular conjugation of As^3+^ with glutathione (GSH) (Kobayashi *et al*., 2005; Hirano, 2020) are included in Fig. 6 for further understanding of the role of As^3+^ on PML proteins.

**Fig. 6.**
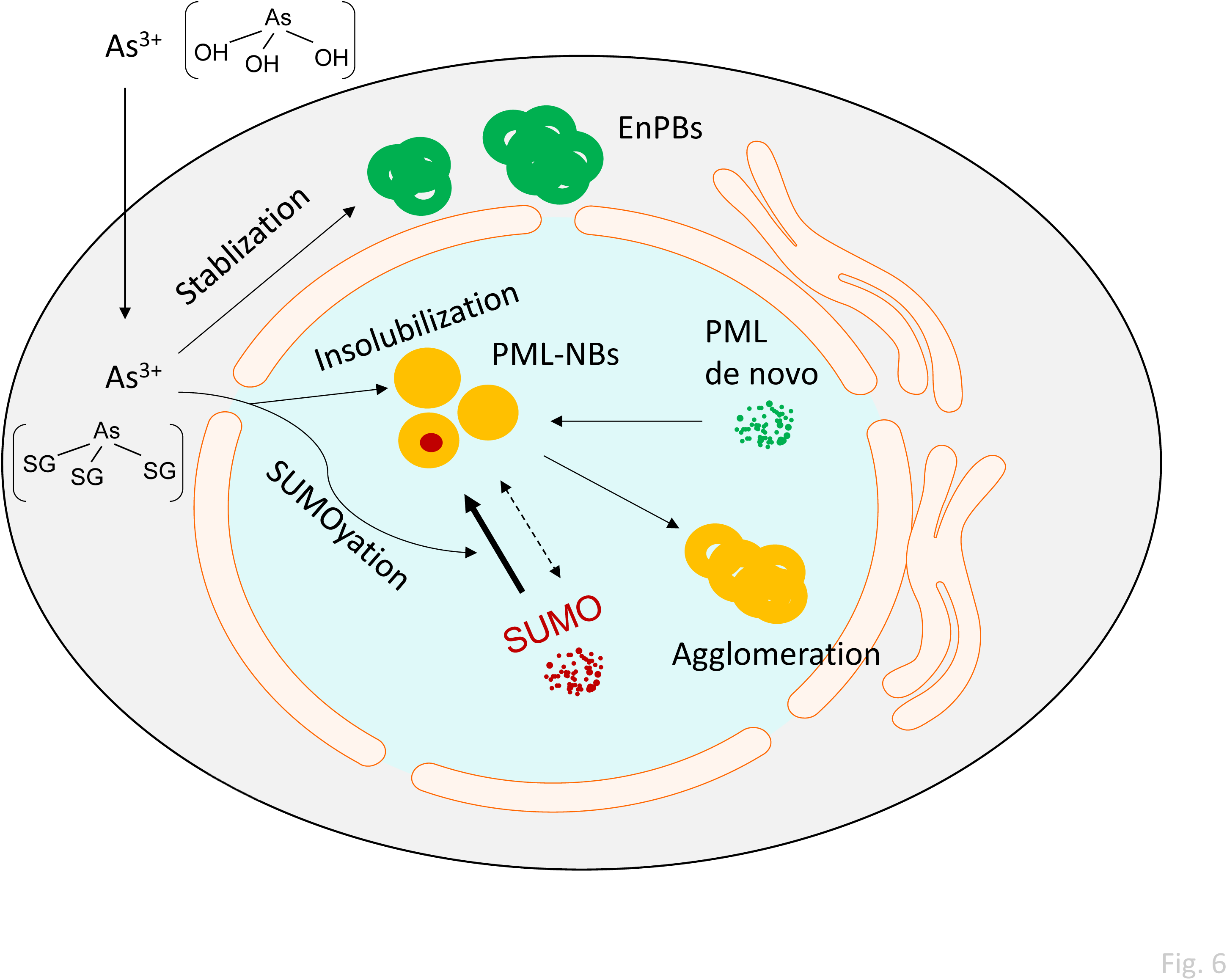
A schematic presentation for the effects of As^3+^ on intranuclear and extranuclear PML. The stability of PML-NBs is regulated by rapid SUMOylation and deSUMOylation on PML proteins, and As^3+^ shifts the equilibrium toward SUMOylation as As^3+^ transforms PML from the soluble to insoluble forms. As^3+^ also stabilizes extranuclear PML (EnPBs) which are reminiscence of PML-NBs at mitosis. See also the text for cellular uptake of As^3+^ and conjugation of As^3+^ with glutathione (GSH).

### Regulation of the number and size of PML-NBs

It is proposed that the initial step for PML-NB formation is a PML dimer generation via a hydrophobic interaction of RING domain (L73 RING dimer) {Wang, 2018 #66}. This process is followed by oligomerization through hydrophobic interaction of B-box and coiled-coil domains {Li, 2019 #122}. Further phase separation occurs through C-terminal disordered region (Silonov *et al*., 2023). It has been shown that L240R, L247R, and I279R mutations in the CC domain impaired PML-NB formation and ATO (2 μM, 1h) partially restored PML-NBs disrupted by these mutations in HeLa cells (Tan *et al*., 2024). It was reported that the C-terminal region of PML-II and PML-V can form nuclear bodies and that of other PML isoforms may be incorporated into PML-NBs independent of their N-terminal regions (Geng *et al*., 2012). As the RING domain, B-box, and coiled-coil in the tripartite motif (TRIM/RBCC) of PML play a critical role in the assembly of PML-NBs, it is reasonable to conclude that various interactions among PML protein isoforms lead to the NB formation (Silonov *et al*., 2023). Our present study suggests that rapid SUMOylation and deSUMOylation of PML regulate the biogenesis of PML-NBs, which in turn determine the number and size of PML-NBs.

The notion that the SUMO-SIM interaction leads to regulation of PML-NBs (Banani *et al*., 2016) may not represent the whole PML-NB processing scheme, because PML-VI which does not has SIM and a PML mutant (3KR) can forms PML-NBs similarly (Brand *et al*., 2010; Hoischen *et al*., 2018; Hirano and Udagawa, 2022a). Although the SUMO-SIM interaction is not requisite for the formation of PML-NBs, the interaction has a potency to change the shape of PML-NBs. It was reported that GFP-tagged wild PML-I to -VI, GFP-PML-I, III, IV, and -V formed toroidal PML-NBs, and GFP-PML-II and VI generated aggregated structures in HeLa cells. PML-I and PML-IV, but not PML-III and PML-V, lost the hollowness in PML-NBs by mutation of SIM, and an addition of SIM to PML-VI produced toroidal PML-NBs (Li *et al*., 2017). We reported that wild PML-VI formed toroidal PML-NBs irrespective of the size and the hollowness remained in the perinuclear cytoplasm or EnPBs in CHO-K1 cells (Hirano and Udagawa, 2022a). Collectively, The SUMO-SIM interaction is not prerequisite for phase separation of PML in the nuclei, and the number, size, and shape of PML-NBs are affected by intermolecular association including SUMO-SIM interaction. The N-terminal region of SUMO1 may play an important role in the formation of PML-NBs in a paralogue-specific manner (Lussier-Price *et al*., 2022).

### Effects of As^3+^ on the assembly of PML-NBs

Most PML proteins engaged in PML-NBs were soluble in cold RIPA in untreated cells and insoluble in As^3+^-exposed cells, suggesting that they are liquid-like in untreated cells and more solid-like after exposure to As^3+^. As^3+^ probably changes the conformation of PML via binding to cysteine residues of RBCC which leads to ”rigidness” of PML-NBs (Hyman *et al*., 2014). The “rigidness” might be related to the shape of phase-separated biocondensates. Cytosolic FUS coalesces with stress granules which are formed by heat- and As^3+^-induced liquid-liquid phase separation (LLPS). Mutation of FUS, which are often observed in ALS patients, enhances the liquid-to-solid transition converting spherical FUS droplets to “sea urchin-like” or “starburst-like” aggregated forms (Patel *et al*., 2015). We reported that PML-NBs agglomerate after prolonged exposure to As^3+^ ans Sb^3+^ (Hirano and Udagawa, 2022a). In the presnt study the agglomeration of PML-NBs were observed after 19 h-exposure to As^3+^ in the prensence of ML792 for the last 18 h (Fig. 4(A)). The binding of As^3+^ to cysteine residues of PML causes confomational change of PML-NB proteins (Zhang *et al*., 2010), which may lead to the agglomeration of PML-NBs as well as PML oligomerization.

The number of PML-NBs increases during G1 and G2 interphase arrests, and treatment with As^3+^ decreased the number of PML-NBs (Baba *et al*., 2025). The present study clearly shows that exposure to As^3+^ for 24 h to 72 h decreased PML-NBs (Figs 1 and 3). It is proposed that PML-NBs function as a nuclear SUMOylation hotspots because both SUMOylation machinery and de-SUMOylation enzymes reside in PML-NBs, and more than a half of PML partner proteins in PML-NBs contain negatively charged amino acid-dependent SUMOylation motif (NDSM) (Van Damme *et al*., 2010). Since PML-NBs interfere with the SUMOylation and deSUMOylation cycles of DNA replication factors (Berchtold *et al*., 2018), the prolonged exposure to As^3+^ may change the cell replication dynamics by regulating availability of SUMO molecules.

### Effects of As^3+^ on the stability of EnPBs

At the prophase-to-prometaphase transition, PML-NBs move rapidly just before nuclear envelop breakdown (Chen *et al*., 2008), which leads to formation of MAPPs and CyNPs. These extranuclear PMLs are either degraded by UPS or recycled for PML-NBs (Jul-Larsen *et al*., 2009; Lang *et al*., 2012). However, our present study showed that EnPBs were stable in the presence of As^3+^ (Fig. 5(A)). The stability of EnPBs may depend on PML isoforms, expression levels of PML, and cell types. When cells were exposed to As^3+^ for 72 h, Most PML were engaged in EnPBs, and PML proteins in EnPBs were much less SUMOylated than those in PML-NBs. SUMOylation hardly occurs on PML molecules in EnPBs as the SUMOylation machinery is located in nuclei (Van Damme *et al*., 2010). Western blotting and fluorescent micrographs indicated that PML proteins in EnPBs were covalently modified by a prolonged exposure to As^3+^ generating low electromobility products. (Figs. 3 and 5). Nuclear pore proteins may play a role in the As^3+^-induced formation of the low electromobility products in EnPBs (Lang *et al*., 2012). A recent proteomic study using *S. cerevisiae* shows that As^3+^-binding proteins are enriched in nucleocytoplasmic transport proteins such as importins, exportins, and nuclear pore proteins, and nuclear import of NLS-containing cargos is disrupted in As^3+^-exposed cells (Lorentzon *et al*., 2025). Importin 90 has been identified as a As^3+^-binding protein in a mammalian cell line MCF-7 (Zhang *et al*., 2007). However, the mechanism of As^3+^-induced modification of PML proteins in EnPBs remains to be answered.

In conclusion, the soluble-to-insoluble shift is an initial and important step for the action of As^3+^ on PML proteins and PML-NBs. The continuous and rapid SUMOylation and deSUMOylation cycle may play a role in intranuclear dynamics of PML-NBs. As^3+^ stabilizes EnPBs where PML proteins are much less SUMOylated. The stabilization of EnPBs by prolonged exposure to As^3+^ surely shed light on clinical use of As^3+^, because this effect occurred at a low concentration of As^3+^ and the stability of EnPBs should increase the availability of SUMO molecules in nuclei.

## Supporting information

Uncropped western images

## Acknowledgment

The authors would like to thank Dr. Ayaka K. Udagawa for technical assistance and discussion.

## Author contributions

SH designed the study, performed experiments, and wrote the paper. OU performed experiments and wrote the paper.

