## Supplementary figures for "Biological differences in promyelocytic leukemia (PML) proteins between PML-nuclear bodies (PML-NBs) and extranuclear PML bodies (EnPBs) in arsenite-exposed cells"

**Supplementary figure legends**

**Supplementary Fig. 1., Western blot analysis for PML and SUMO2/3 in U-2OS (A) and HEK293 cells (B).** (A) U-2OS cells were exposed to 3 μM $As^{3+}$ for 4, 24, and 48 h. The cells were further cultured in fresh complete culture medium for 24 or 48 h after 4 h-exposure to 3 μM $As^{3+}$. (B) HEK293 cells were exposed to 3 μM $As^{3+}$ for 2, 4, 24, or 72 h or 0.3 μM $As^{3+}$ for 72 h. A right curly bracket and an open arrowhead indicate PML isotypes and a SUMO monomer, respectively.

**Supplementary Fig. 2., Semi-quantitative analyses of PML-VI, SUMO1 and SUMO2/3 monomers, and SUMOlyated PML-VI in *PML(VI)* U-2OS cells.** The cells were exposed to 3 μM $As^{3+}$ (As) or left untreated (-) for 72 h, and lyzed in ice-cold RIPA buffer. Western blot analysis was performed for PML, DDK (FLAG), SUMO2/3, and SUMO1 in the RIPA-soluble and -insoluble fractions. Closed and open arrowheads indicate PML-VI and SUMO monomers, respectively. Note that PML-VI was expressed with DDK (FLAG) tag in the present mammalian expression system. Data are presented as mean ± SEM (N=3) *, Significantly different from the untreated wells (-).

**Supplementary Fig. 3., Western blot analysis for PML and SUMO2/3 and in CHO-K1 cells stably expressing GFP-PML-VI and wild-type (CHOgPrSWild), K11R mutant (CHOgPrS (K11R)) or C-terminal diglycine-deleted mCherry-tagged SUMO2 (CHOgPrSΔGG) cells.** (A) CHOgPrSWild, CHOgPrS(K11R), or CHOgPrSΔGG cells were exposed to 3 μM $As^{3+}$ (As) or left untreated for 2 h (-). The membrane was stained with Ponseau-S before blocking. (B) Effects of ML792 on SUMOylation of PML in CHOgPrS(K11R), or CHOgPrSΔGG cells. The cells were pre-treated with 20 μM ML792 for 1 h and then exposed 3 μM $As^{3+}$ for 2 h. Closed and open arrowheads indicate PML-VI and SUMO monomers, respectively. An arrow indicates a mCherry-tagged SUMO2. A right square and a right double square bracket indicate GFP-tagged PML-VI SUMOylated with endogenous SUMO2/3 and GFP-tagged PML-VI SUMOylated mainly with mCherry-tagged SUMO2, respectively.

**Supplementary Fig. 4., Immunofluorescence micrographs of CHOgPrSWild cells.** The cells were left untreated or exposed to $Cd^{2+}$, $As^{3+}$, $Cu^{2+}$, or $Sb^{3+}$ at indicated concentrations for 72 h. Scale bar = 20 μm. An arrow indicates an EnPB.

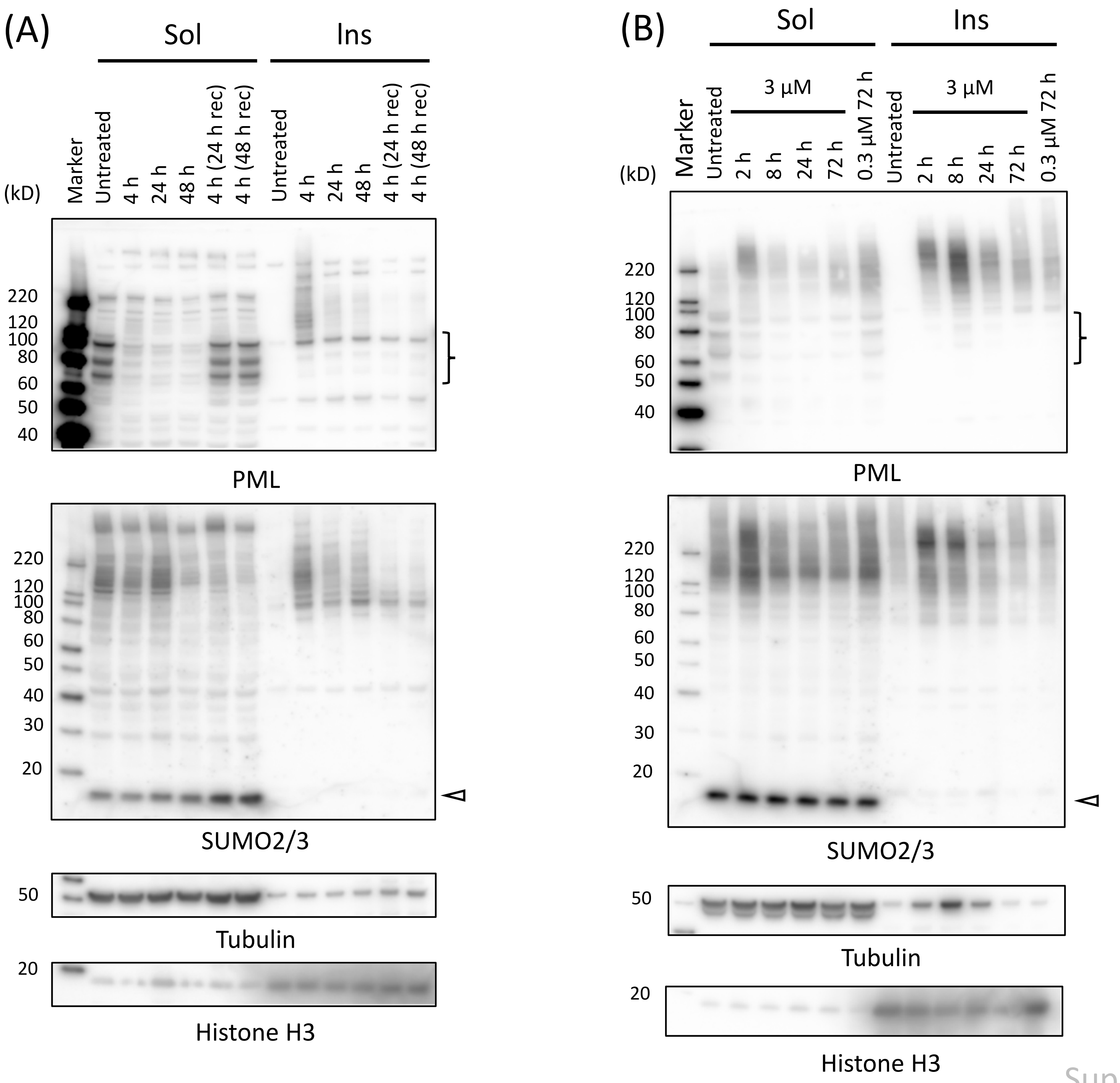


Supplementary Fig. 1

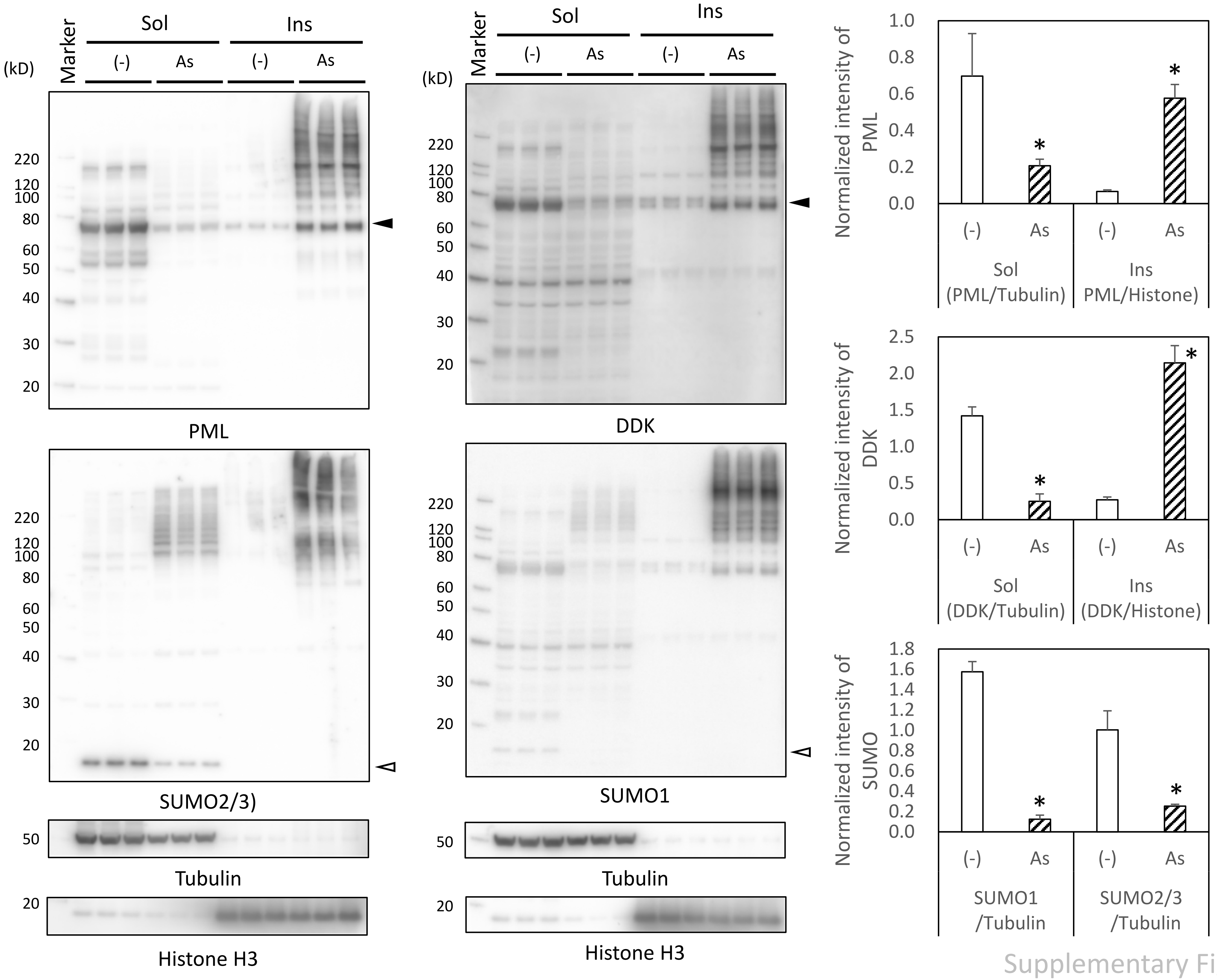


Supplementary Fig. 2

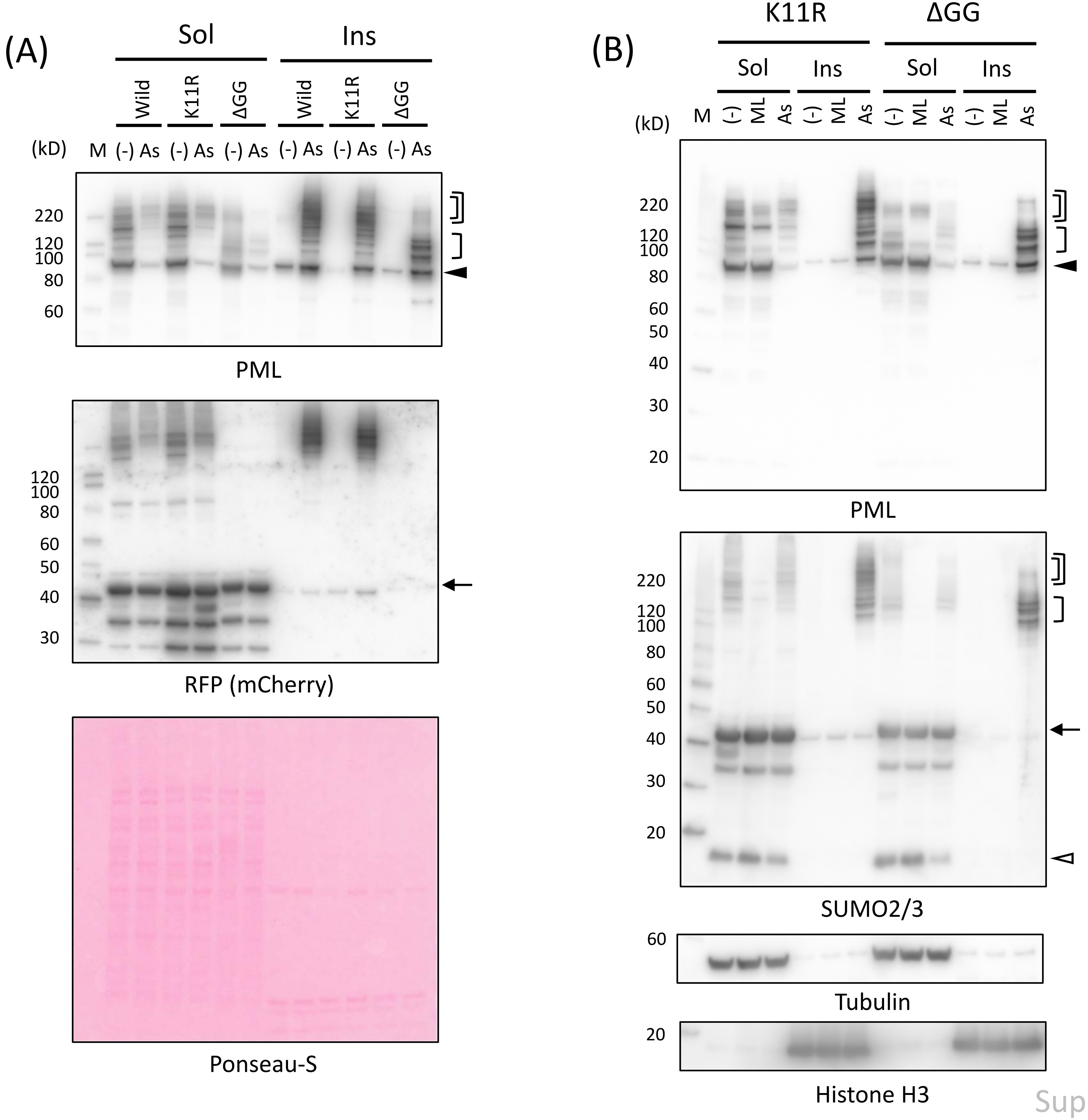


Supplementary Fig. 3

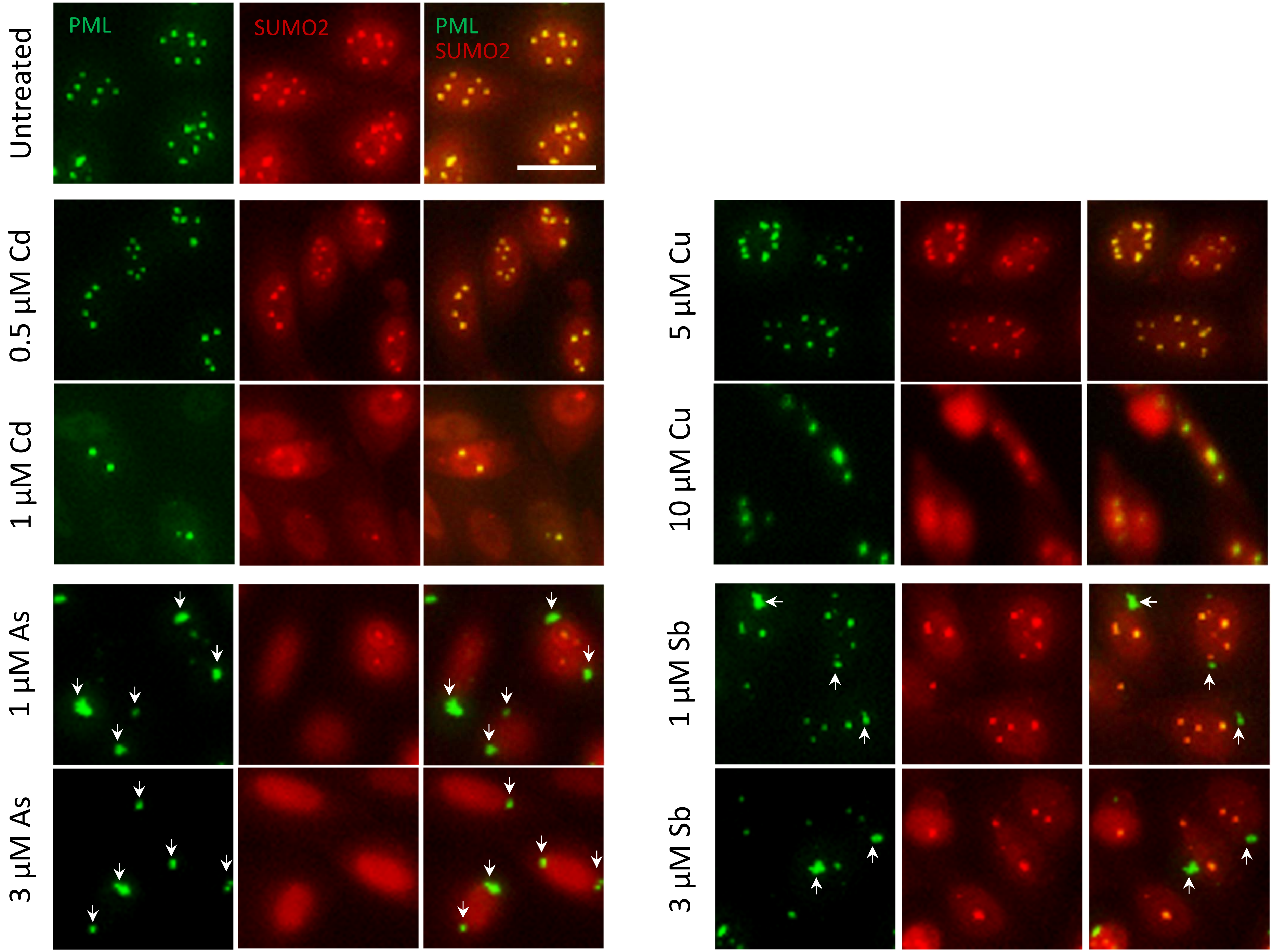


Supplementary Fig. 4
