## Supplementary material for "Biological differences in promyelocytic leukemia (PML) proteins between PML-nuclear bodies (PML-NBs) and extranuclear PML bodies (EnPBs) in arsenite-exposed cells": Source table

Table 1, Information for antibodies and plasmids used in the study.

| Reagent | Name | Source | Identifier | Dilution rate |
|---|---|---|---|---|
| Primary antibody | anti-PML, Rabbit, polyclonal | Bethyl | Cat# A301-167A-3 | 1:1000 for WB, 1:125 for IF |
| | anti-SUMO1 Rabbit, monoclonal | CST, Clone C9H1 | Cat# 4940 | 1:1000 for WB |
| | anti-SUMO2/3, Mouse, monoclonal, IgG2b | MBL, Clone 1E7 | Cat# M114-3 | 1:1000 for WB |
| | anti-ubiquitin Mouse, monoclonal | Cytoskelton, Clone P4D1 | Cat#: AUB01 | 1:1000 for WB |
| | anti-RFP cocktail, Mouse monoclonal IgG2bκand IgG1κ | MBL, Clone 1G9 3G5 (mixed) | Cat# M208-3 | 1:1000 for WB |
| | anti-GFP, Goat, polyclonal | Abcam | Cat# ab5450 | 1:1000 for WB |
| Secondary antibody | goat anti-Rabbit IgG (HRP) | Santa Cruz | Cat#: sc2054 | 1:2500 for WB |
| | goat anti-Mouse IgG (HRP) | Santa Cruz | Cat#: sc2055 | 1:2500 for WB |
| | mouse anti-Goat IgG (HRP) | Santa Cruz | Cat#: sc2354 | 1:2500 for WB |
| | Alexa Fluor™ 488-labeled goat anti-mouse IgG | Invitrogen-Thremo Fisher | Cat#: A31620 | 1:1000 for IF |
| | Alexa Fluor™ 594-labeled goat anti-rabbit IgG | Invitrogen-Thremo Fisher | Cat#: A31632 | 1:1000 for IF |
| Conjugated antibody | anti-α-tubulin (HRP) Rabbit, polyclonal | MBL | Cat# PM054-7 Anti-α-tubulin HRP-DirectT | 1:2000 for WB |
| | anti-histone H3 (HRP) Rabbit, monoclonal, IgG2b | CST, Clone 3H1 | Cat# 12648 HRP-conjugated anti-histone H3 | 1:1000 for WB |
| Plasmid | Myc-DDK-tagged ORF Homo sapiens PML transcript variant 5 | OriGene | RC220236 [NM_033244.3] | |
| | PML doubke nickase plasmid | Santa Cruz | sc400145-NIC | |
| | mCherry-SUMO2 Wild | VectorBuilder, mCherry/mSumo2 | VB210203-1053ufy [NM_133354.2] | |
| | mCherry-SUMO2 K11R mutant | VectorBuilder, mCherry/mSumo2 | VB210203-1267fwb [NM_133354.2] | |
| | mCherry-SUMO2 C-terminus GG deleted mutant | VectorBuilder, mCherry/mSumo2 | VB210203-1259pus [NM_133354.2] | |
