## Supplementary material for "Biological differences in promyelocytic leukemia (PML) proteins between PML-nuclear bodies (PML-NBs) and extranuclear PML bodies (EnPBs) in arsenite-exposed cells": Uncropped western images

Fig. 1B PML

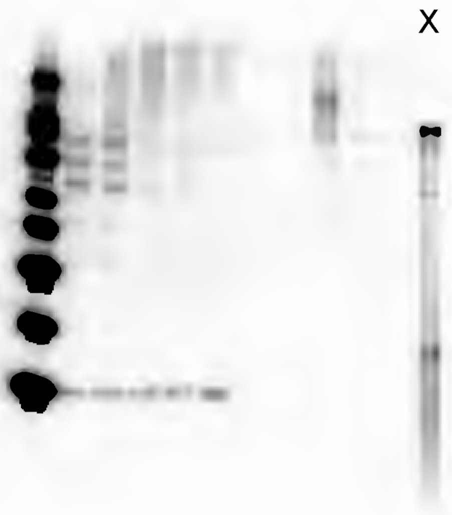

Fig. 1B SUMO23

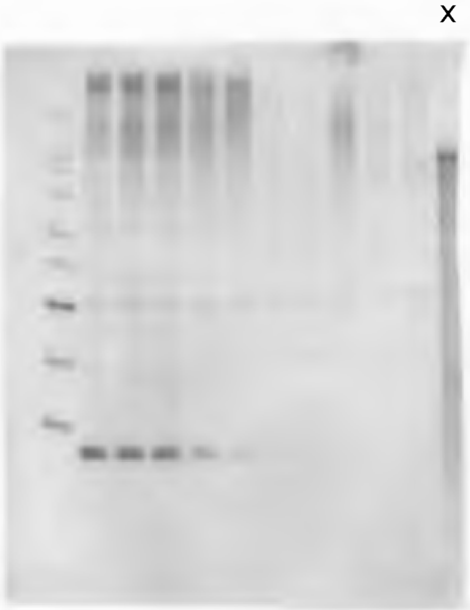

Fig. 1B Tubulin
Histone

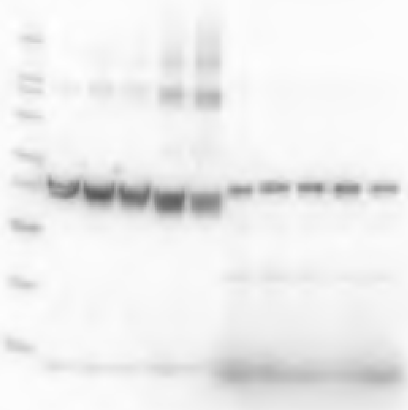

Fig. 1C PML

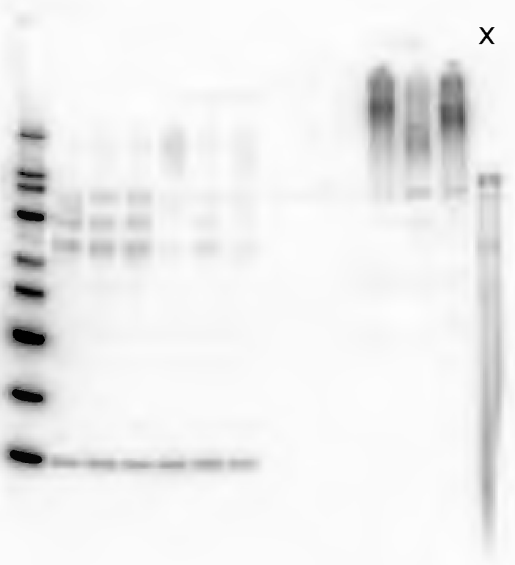

Fig. 1C SUMO23

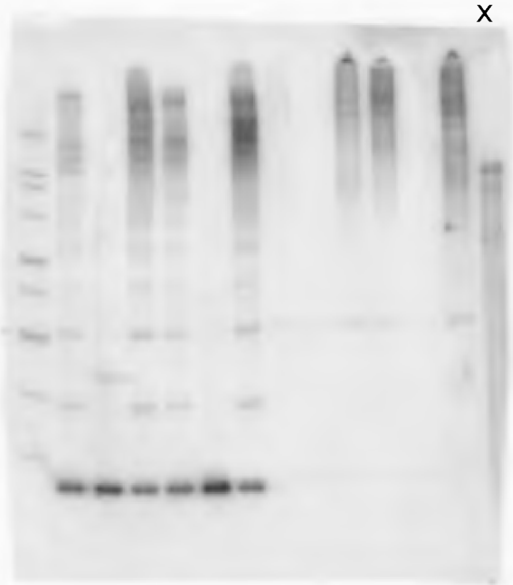

Fig. 1C Tubulin
Histone (left)

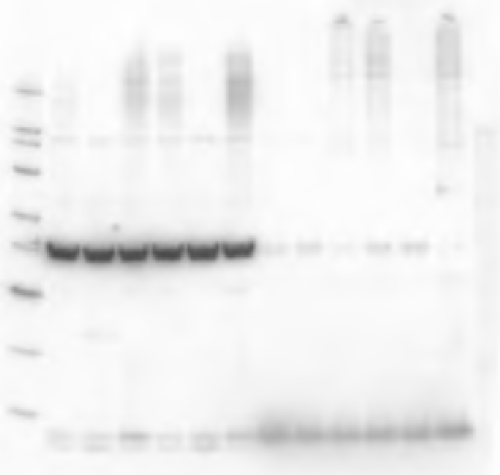

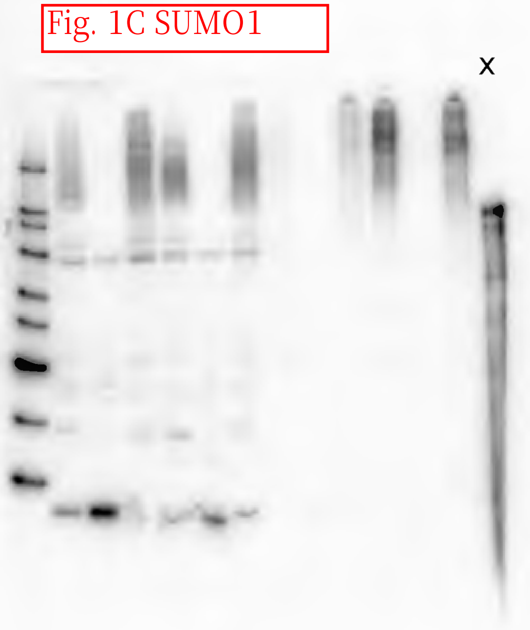
Fig. 1C SUMO1
x

Fig. 1C Ubiquitin

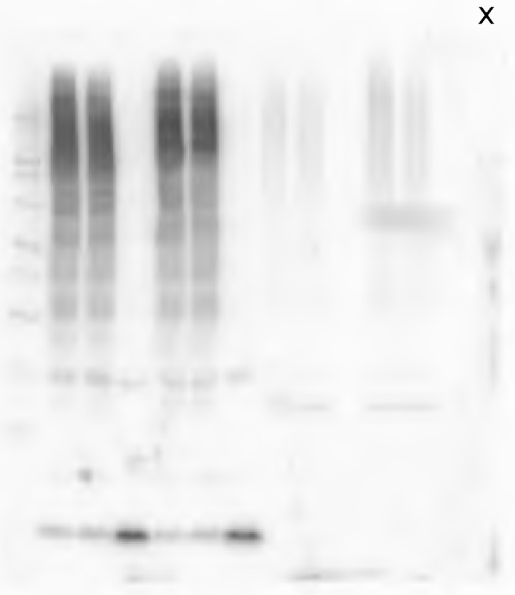

Fig. 1C Tubulin
Histone (right)

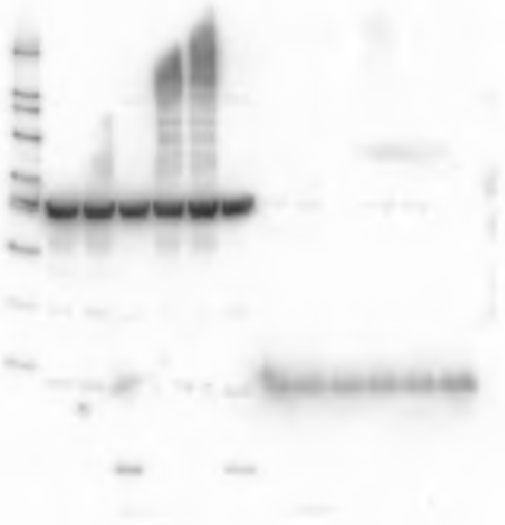

Fig. 1E PML

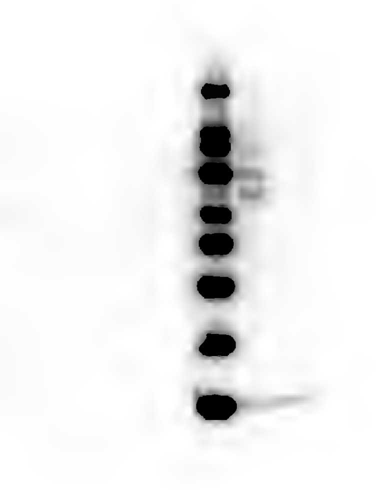

Fig. 1E SUMO23

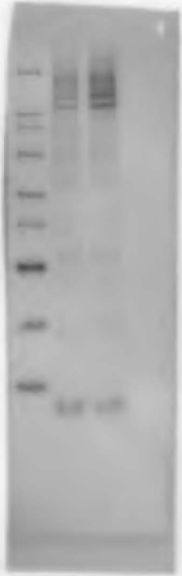

Fig. 1E Tubulin

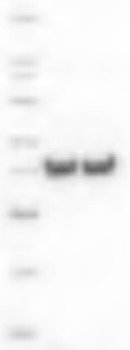

Fig. 2C PML

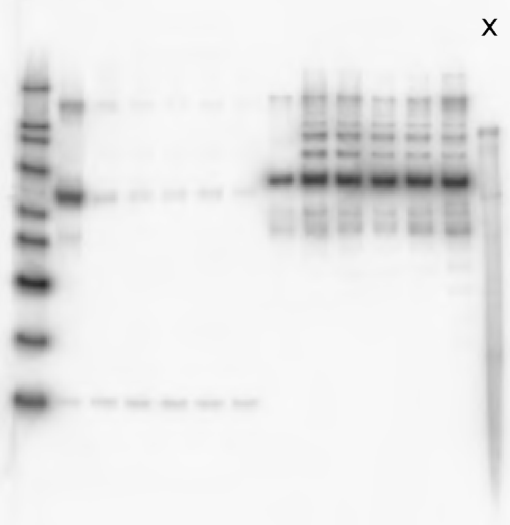

Fig. 2C Tubulin
Histone

x

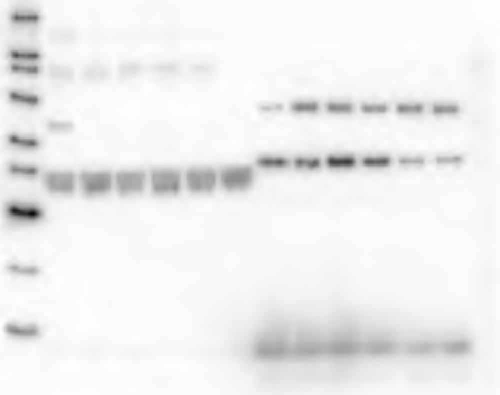

Fig. 2C SUMO23

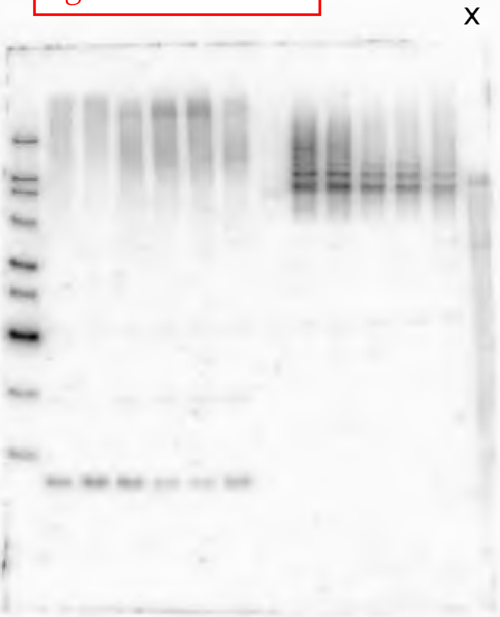

Fig. 3F PML

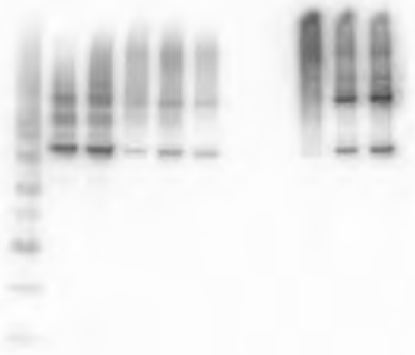

Fig. 3F SUMO23

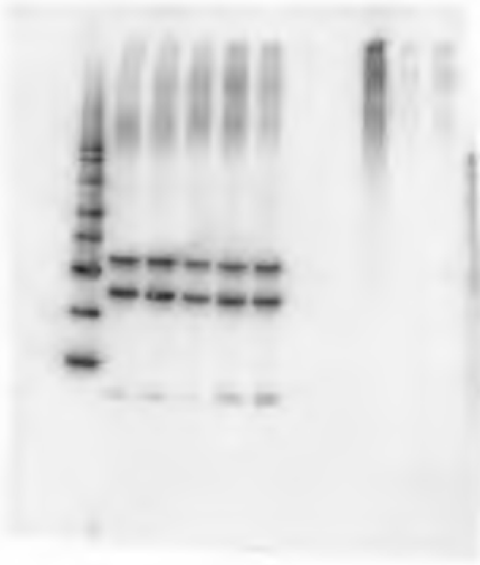

Fig. 3F Tubukin Histone (left)

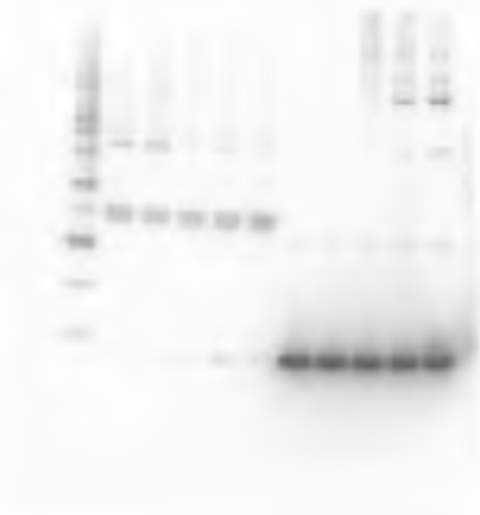

Fig. 3F SUMO1

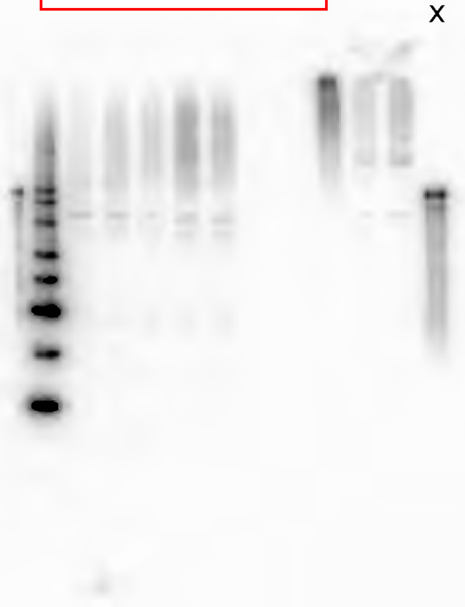

Fig. 3F Ubiquitin

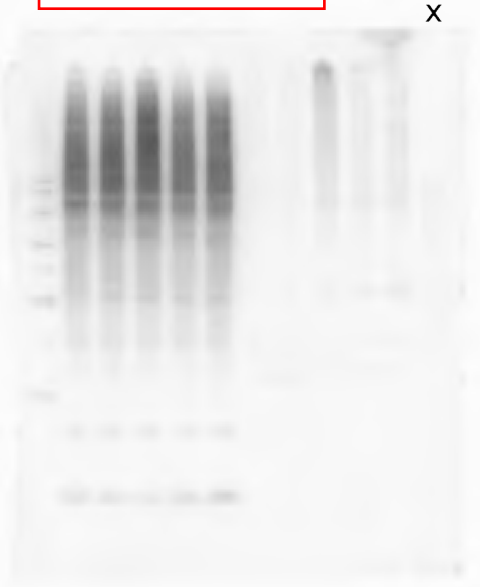

Fig. 3F Tubulin
Histone (right)

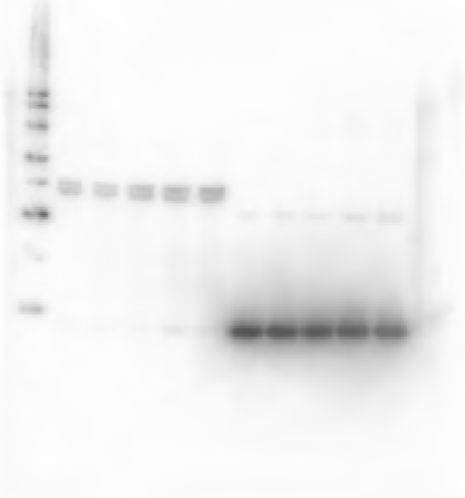

Fig. 4B PML

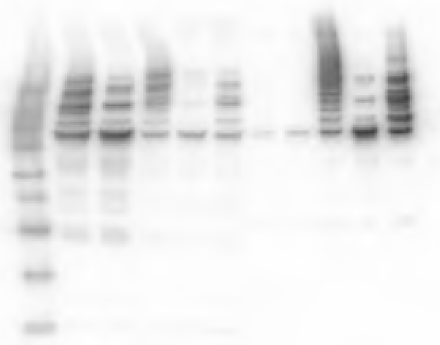

Fig. 4B SUMO23

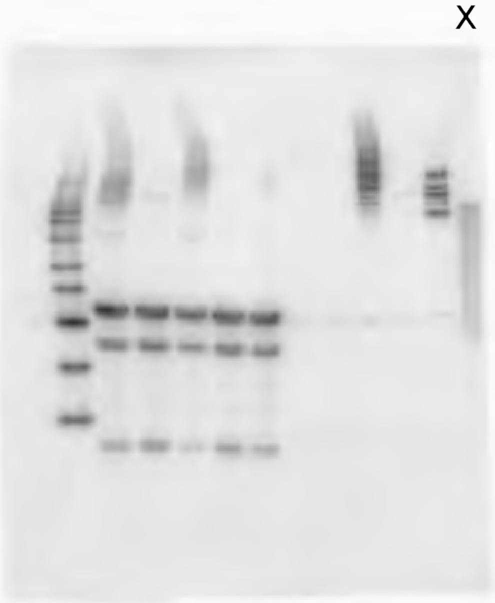

Fig. 4B Tubulin
Histone (left)

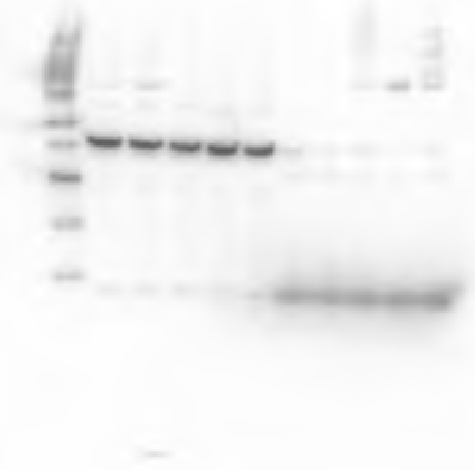

Fig. 4B GFP

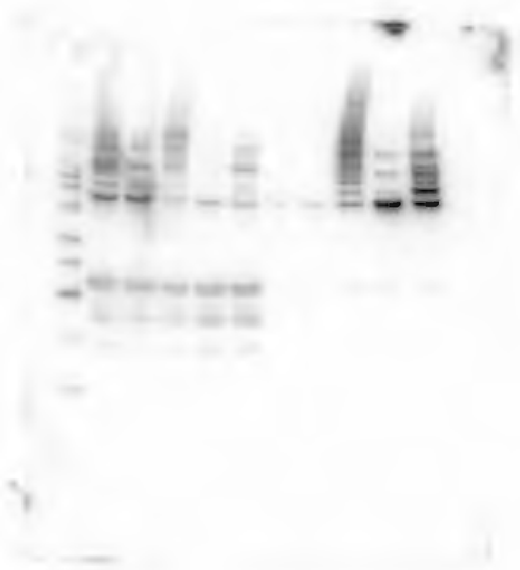

Fig. 4B RFP

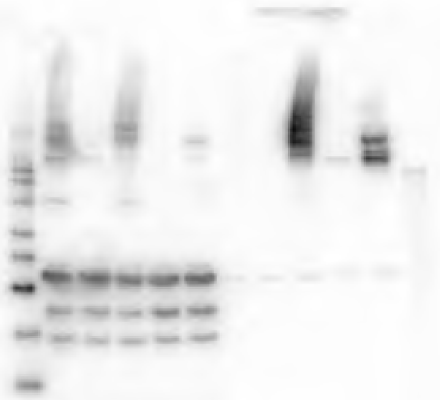

Fig. 4B Tubulin
Histone (right)

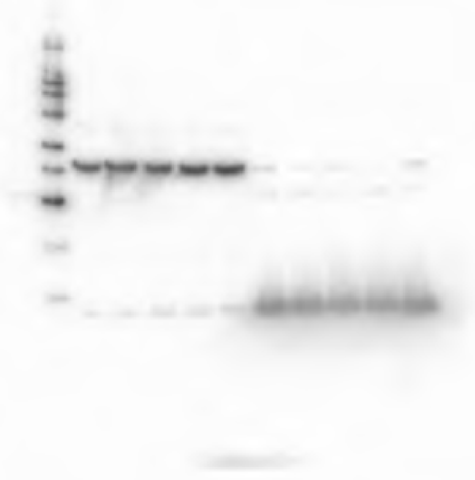

Fig. 5B PML

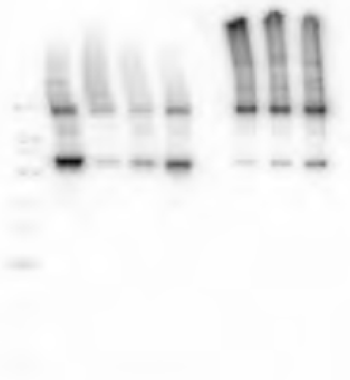

Fig. 5B SUMO23

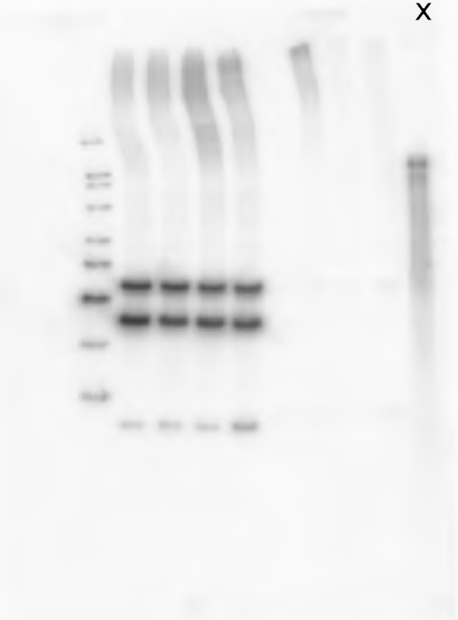

Fig. 5B Tubulin
Histone (left)

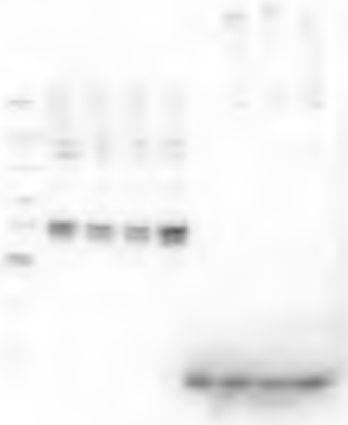

Fig. 5B SUMO1

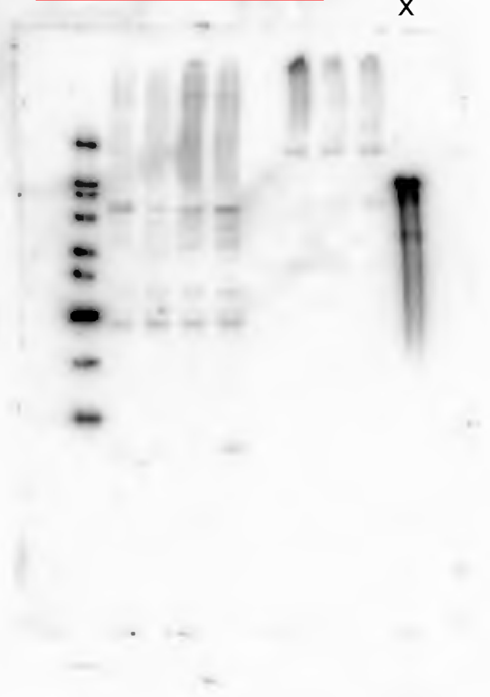

Fig. 5B Ubiquitin

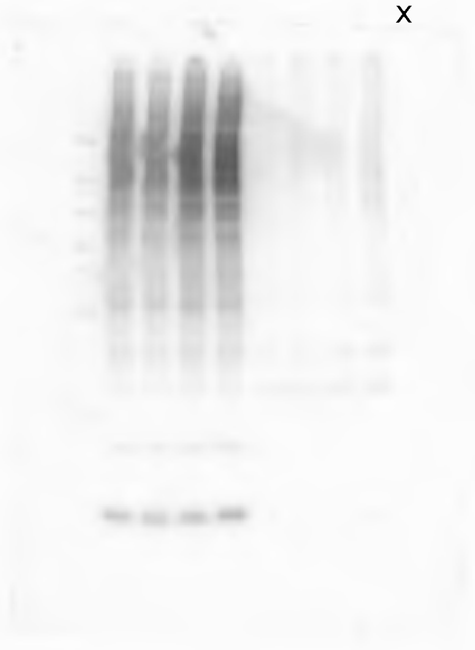

Fig. 5B Tubulin
Histone (right)

x

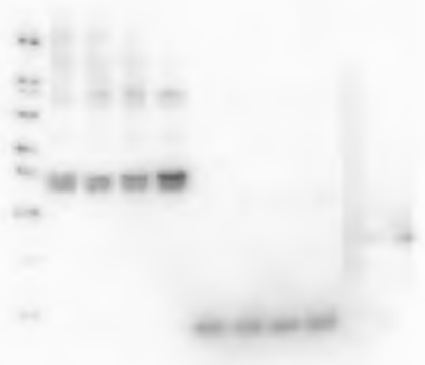

S.Fig. 1A PML

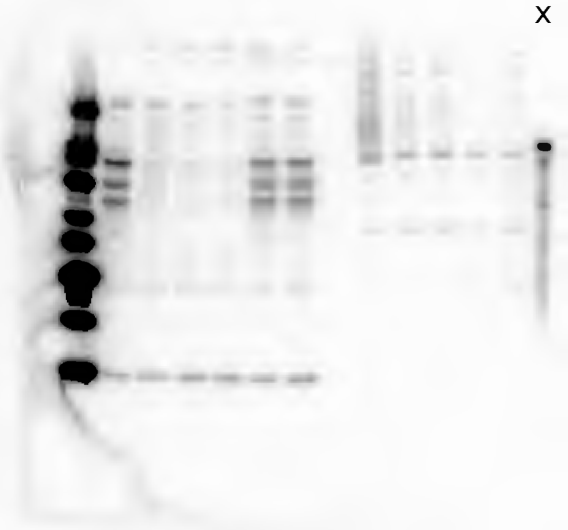

S.Fig. 1A SUMO23

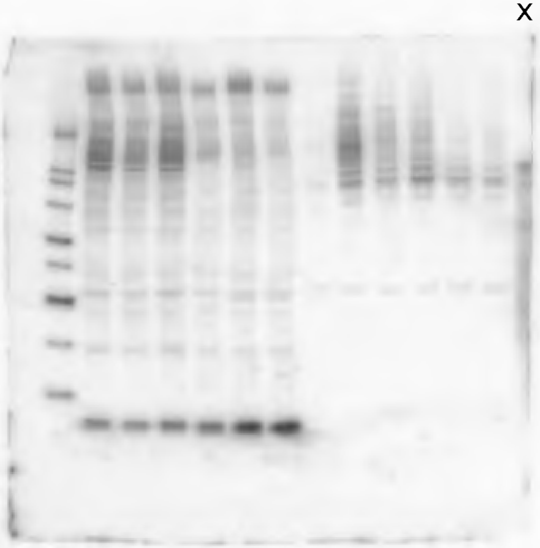

S.Fig. 1A Tubulin Histone

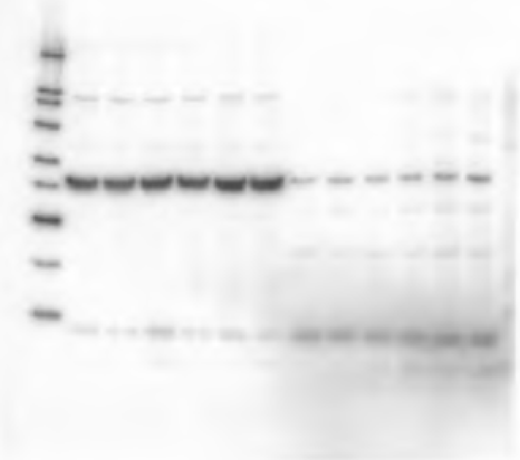

S.Fig. 1B PML

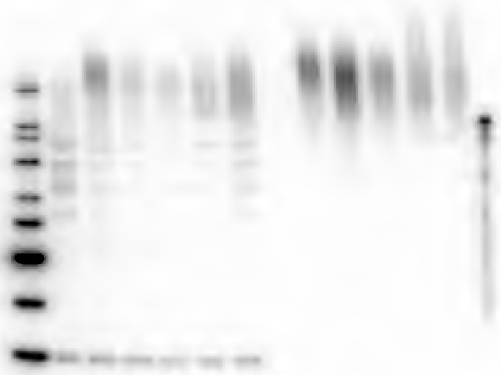

S.Fig. 1B SUMO23

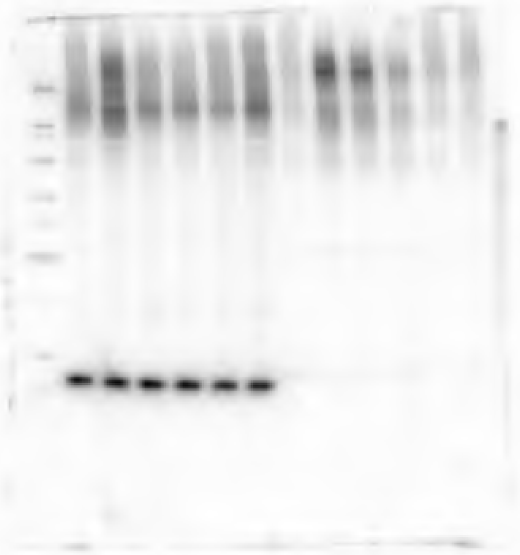

S.Fig. 1B Tubulin Histone

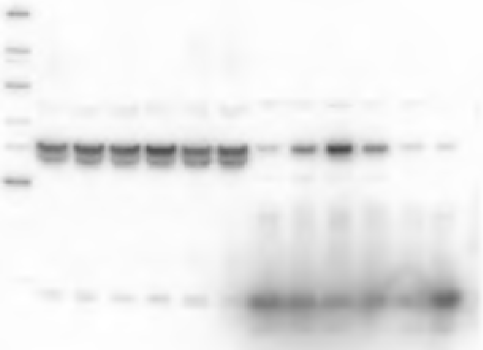

S.Fig.2 PML

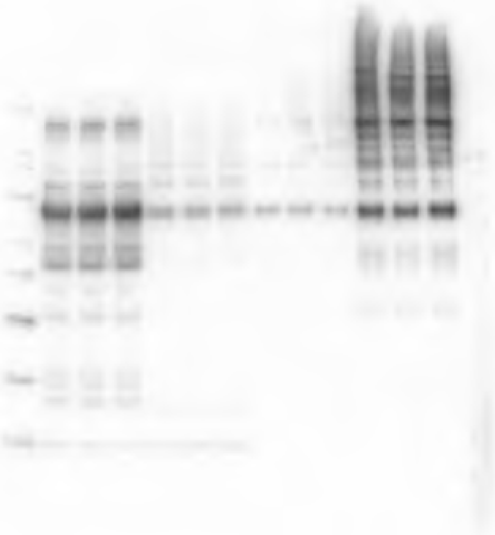

S.Fig.2 SUMO23

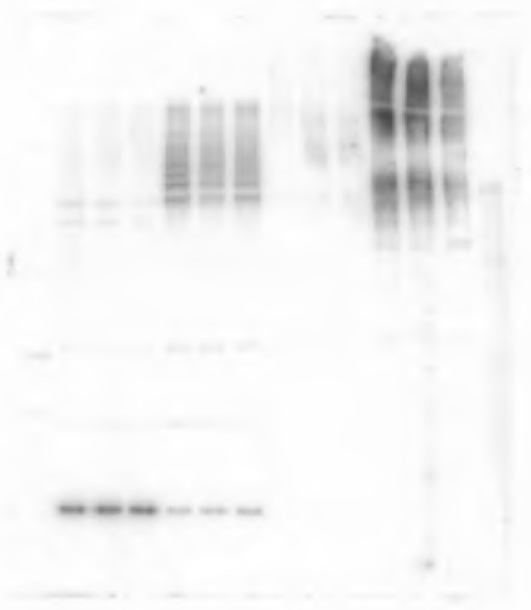

S.Fig.2 Tubulin
Histone (left)

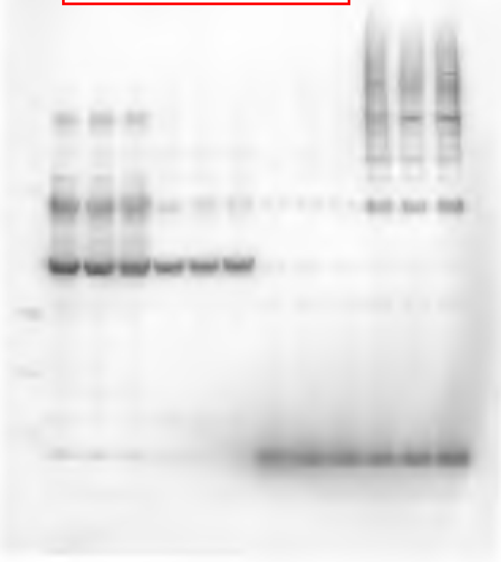

S.Fig.2 DDK

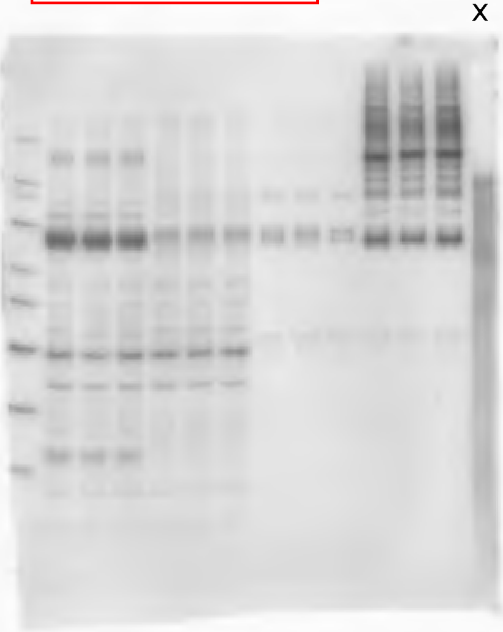

S.Fig.2 SUMO1

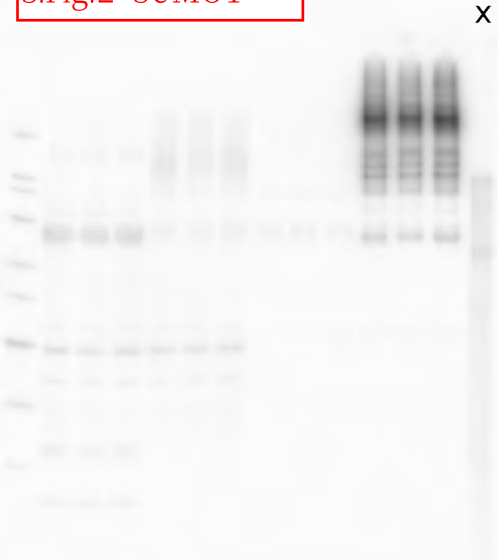

S.Fig.2 Tubulin
Histone (right)

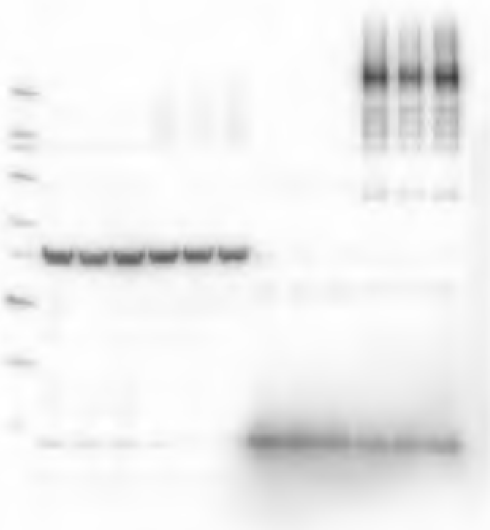

S.Fig. 3A PML

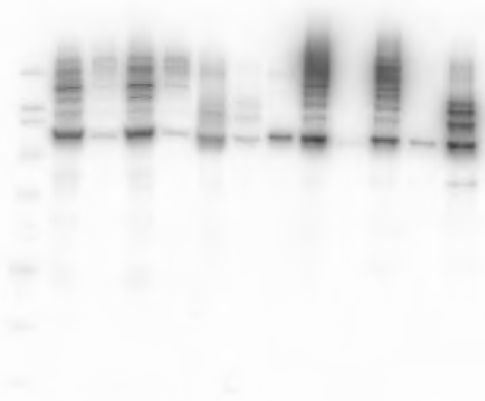

S.Fig. 3A RFP

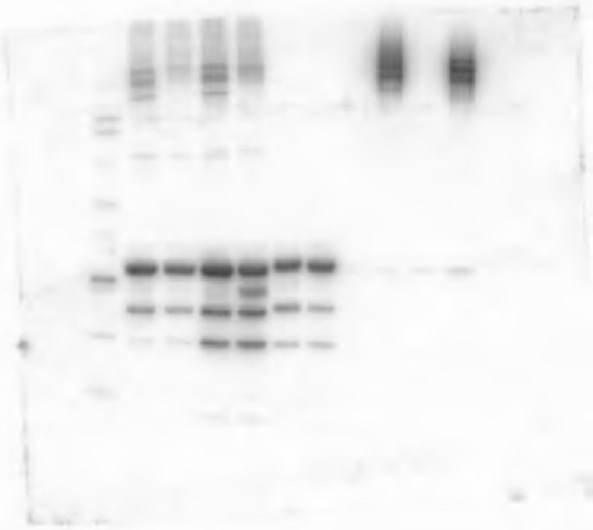

S.Fig. 3B PML

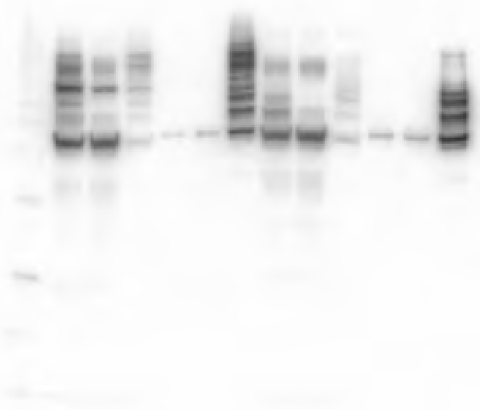

S.Fig. 3B SUMO23

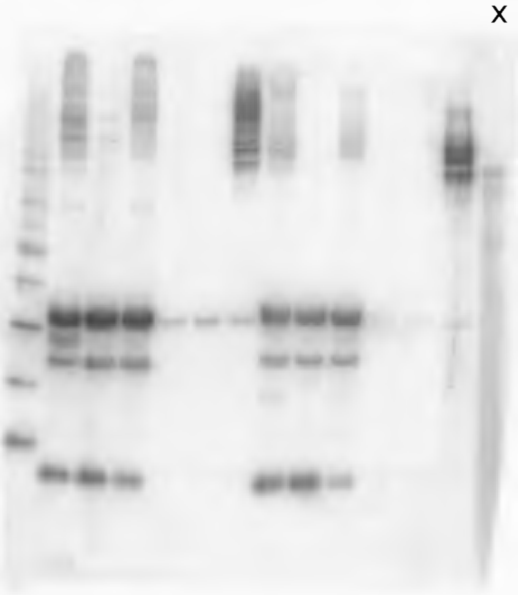

S.Fig. 3B Tubulin
Histone

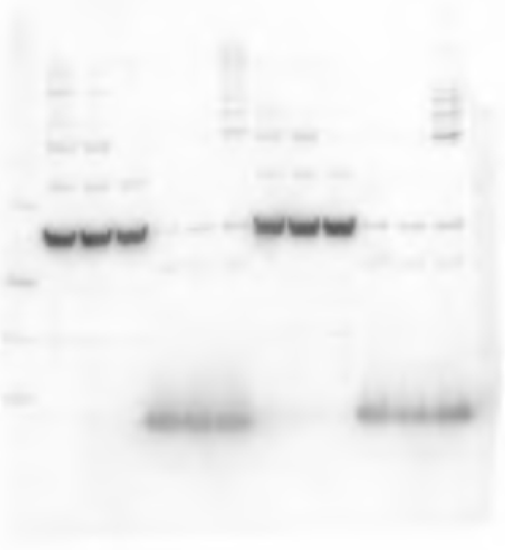
